# Zinc Finger MYND-Type Containing 8 (ZMYND8) is epigenetically regulated in mutant Isocitrate Dehydrogenase 1 (IDH1) glioma to promote radioresistance

**DOI:** 10.1101/2022.06.10.495694

**Authors:** Stephen V. Carney, Kaushik Banerjee, Anzar Mujeeb, Brandon Zhu, Santiago Haase, Maria L. Varela, Padma Kadiyala, Claire Tronrud, Ziwen Zhu, Devarshi Mukherji, Preethi Gorla, Rebecca Tagett, Felipe J. Núñez, Maowu Luo, Weibo Lou, Mats Ljungman, Yayuan Liu, Ziyun Xia, Anna Schwendeman, Tingting Qin, Maureen A. Sartor, Joseph Costello, Daniel P. Cahill, Pedro R. Lowenstein, Maria G. Castro

## Abstract

Mutant isocitrate dehydrogenase 1 (mIDH1) alters the epigenetic regulation of chromatin, leading to a hypermethylation phenotype in adult glioma. Establishment of glioma-specific methylation patterns by mIDH1 reprogramming drives oncogenic features of cancer metabolism, stemness and therapeutic resistance. This work focuses on identifying gene targets epigenetically dysregulated by mIDH1. Treatment of glioma cells with mIDH1 specific inhibitor AGI-5198, upregulated gene networks involved in replication stress. Specifically, we found that the expression of ZMYND8, a regulator of DNA damage response was decreased in three patient-derived glioma cell cultures (GCC) after treatment with AGI-5198. ZMYND8 functions as a chromatin reader to modulate enhancer activity and recruit DNA repair machinery. Knockdown of ZMYND8 expression sensitized mIDH1 GCCs to radiotherapy marked by decreased cellular viability. Following IR, mIDH1 glioma cells with ZMYND8 knockout (KO) exhibit significant phosphorylation of ATM and sustained γH2AX activation. ZMYND8 KO mIDH1 GCCs were further responsive to IR when treated with either BRD4 or HDAC inhibitors. The accumulation of DNA damage in ZMYND8 KO mIDH1 GCCs promoted the phosphorylation of cell cycle checkpoint proteins Chk1 and Chk2. In addition, the recruitment of ZMYND8 to sites of DNA damage has been shown to be PARP- dependent. PARP inhibition further enhanced the efficacy of radiotherapy in ZMYND8 KO mIDH1 glioma cells. These findings indicate the impact of ZMYND8 in the maintenance of genomic integrity and repair of IR-induced DNA damage in mIDH1 glioma.

**Translational Relevance:** Our understanding of radioresistance mechanisms in patient-derived glioma cell cultures (GCC) that endogenously express mIDH1-R132H are limited. We have uncovered a novel gene target Zinc Finger MYND-Type Containing 8 (ZMYND8) that is downregulated following treatment of human mIDH1 GCCs with mIDH1 specific inhibitors. We demonstrate that suppression of ZMYND8 expression by shRNA knockdown or genetic knockout reduces the cellular viability of mIDH1 GCCs to ionizing radiation (IR). Our findings reveal an epigenetic vulnerability of mIDH1 GCCs to ZMYND8 knockout (KO) which results in impaired resolution of IR-induced DNA damage and induction of cell cycle arrest. Additionally, ZMYND8 KO mIDH1 GCCs display increased radiosensitivity to inhibition of epigenetic regulators BRD4, HDAC, and PARP which could be mediated by enhanced replicative stress.

## Introduction

IDH-mutant gliomas account for roughly 80% of low grade gliomas. (1) The integration of histological classification with genomic analysis of concurrent mutations has defined the subtypes of IDH-mutant glioma as either astrocytoma (TP53 and ATRX inactivating mutations), oligodendroglioma (TERT promoter amplification, 1p19q co-deletion), or recurrent glioblastoma (CDKN2A loss, CDK4 amplification). (2) The low grade mIDH1 tumors are slower growing, but highly infiltrative and arise predominantly within the frontal lobe. (3) Patients harboring mutations in either IDH1 or IDH2 have a favorable prognosis as compared with high-grade wildtype-IDH glioma. (4) Despite this finding, many IDH-mutant glioma patients treated with radiotherapy develop tumor progression and eventually succumb to the disease. (5) The identification of molecular mechanisms that drive therapeutic response to IR in IDH1 glioma remains unexplored. In this respect, a recent report discusses recommendations for the implementation of radiation therapy taking into account the WHO 2021 classification of adult-type mutant IDH1 diffuse gliomas, which takes into account both molecular and histopathological features. (6)

The characteristic heterozygous IDH1-R132H mutation produces an oncometablite 2- hydroxyglutarate (2HG) that impairs nuclear enzymes Jumonji-family of histone demethylases and TET2 methylcytosine dioxygenase inducing a IDH-specific hypermethylation phenotype. (7, 8) By reversing the epigenetic reprogramming elicited by IDH-mutations, we can elucidate how cellular pathways altered by mIDH1 support survival of tumor cells following radiation. In this study, we sought out to uncover gene targets that displayed alterations in chromatin regulation and gene expression following mIDH1 inhibition, and in turn discern their contribution to radioresistance. We have previously demonstrated that murine tumor neurospheres (NS) harboring mIDH1^R132H^ mutation display differential enrichment of H3K4me3 methylation at genomic regions associated with master regulators of the DNA Damage Response (DDR). (9) When we inhibited the DNA damage sensor ATM that initiates homologous recombination (HR) pathway or its downstream targets cell cycle checkpoint kinases 1 (Chk1) or 2 (Chk2), we observed enhanced radiosensitization of mIDH1-expressing GCCs. We also showed that pharmacological inhibition of mIDH1-R132H mutation in murine NS reduced mitotic cell proliferation and enhanced the release of Damage Associated Molecular Patterns. (10) Additionally, administering a mIDH1 specific inhibitor (AGI-5198) in combination with ionizing radiation (IR) to mIDH1 tumor-bearing mice promoted tumor regression and the development of immunological memory. (10)

We targeted mIDH1 using AGI-5198, which has been shown to decrease the production of 2HG and reverse histone hypermethylation. (11) Herein, we evaluated changes in gene expression and epigenetic regulation in a radioresistant mIDH1 patient-derived GCC following mIDH1 inhibition. We identified the significant downregulation of the chromatin reader protein, Zinc Finger MYND-Type Containing 8 (ZMYND8, also referred to as RACK7 and PRKCBP1). The connection between ZMYND8 and DNA repair was proposed based on its recruitment to nuclear sites of laser induced DNA damage. (12) The presence of a bromodomain (BRD)-containing motif encoded within ZMYND8, was shown to recognize acetylated lysine residues, specifically H3K14ac and H4K16ac that are present on post-translational modified (PTM) histones. (13) Recent high resolution crystal structure analysis of the ZMYND8 protein, explored the putative chromatin reader function of adjacent domains to bind specific histone marks. The plant homeodomain (PHD) recognizes singly methylated lysines (H3K4me1), while the hydrophobic Pro-Trp-Trp-Pro (PWWP) domain recognizes H3K36me2. (13) These effector domains support early findings that ZMYND8 accumulated at actively transcribed regions where double stranded breaks (DSB) occurred. (12)

During DNA repair, there are dynamic changes to chromatin organization, which allows for the recruitment of DDR proteins. Sites of DNA damage display reductions in active histone mark H3K4me3 signal that co-localizes with hallmark DNA damage markers like γ-H2AX. (14) The recruitment of ZMYND8 to regions of DNA damage is dependent on H3K4me2/3 demethylase KDM5A, proposing a role of transcriptional repression in DNA repair mediated by HR mechanisms. (14) ZMYND8 interacts with several subunits of the Nucleosome Remodeling and Histone Deacetylase (NuRD) complex: Chromodomain Helicase DNA Binding Protein 4 (CHD4), histone deacetylases 1 and 2 (HDAC1/2) and GATA Zinc Finger Domain Containing 2A (GATAD2A) to facilitate HR-dependent DNA repair. (15) This process is shown to be regulated by Poly ADP-ribose polymerase (PARP) dependent recruitment to regions of DNA damage to facilitate a cascade of transcriptional repression. (15) Previous work has shown that ZMYND8 associates with activator complexes like pTEF-b and BRD4 to regulate gene expression at enhancers and active gene regions. (16, 17) Within the realm of tumor biology, ZMYND8 has been linked to the regulation of cancer-specific programs in colorectal, prostate, breast, H3.3G34R mutant glioma, acute myeloid leukemia, renal cell carcinoma, non-small cell lung cancer (NSCLC), nuclear protein in testis (NUT) carcinoma and hepatocellular carcinoma. (18–26)

Herein, we explore resistance mechanisms reinforced by IDH1 reprogramming that allows tumor cells to survive radiotherapy. We have previously shown, mIDH1 GCCs are more resistant to radiotherapy and that inhibition of mIDH1 using AGI-5198 enhances cellular death following radiation. (9, 10) In this study, we utilize three mIDH1 GCCs and our genetically engineered tumor NS that endogenously express IDH1-R132H mutation in the context of ATRX and TP53 loss (NPAI). When we reverse IDH1 reprogramming by treating a human mIDH1 GCC with AGI- 5198, we observe significant reduction in ZMYND8 expression and induction of replication stress genes. Mouse mIDH1 glioma cells express higher ZMYND8 transcripts and display elevated H3K4me3 methylation at the promoter region of ZMYND8 when compared to wildtype IDH1 (wtIDH1) mouse glioma cells. Knocking out ZMYND8 in two mIDH1 patient derived GCC and our genetically engineered mIDH1 mouse tumor neurospheres sensitizes the glioma cells to radiation. Our findings demonstrate that loss of ZMYND8 in mIDH1 GCCs induces greater DNA damage based on higher levels of ATM and γH2AX phosphorylation. This decrease in cellular viability of ZMYND8 KO mIDH1 GCC could be the result of cell cycle arrest induced by p-Chk1 activation. To date, ZMYND8 has been implicated in driving cancer-specific programs mediated by its function at enhancers and through its role in PARP-dependent transcriptional repression. Our data supports the clinical evaluation of PARP, BRD4, or HDAC inhibition to interfere with ZMYND8 mediated repair mechanisms and improve the therapeutic efficacy of irradiation.

## Materials and Methods

### Cell Culture

Human SF10602 cells were grown in NeuroCult NS-A Basal Medium supplemented with 100units/mL antibiotic-antimycotic, 10mL B-27 without vitamin A, 5mL N-2, 100µg/ml Normocin, 20ng/mL FGF, 20ng/mL EGF, and 20ng/mL PDGF-AA. Human MGG119 and LC1035 GCCs were grown in Neurobasal media supplemented with 100units/mL antibiotic-antimycotic,10mL B-27, N-2, 100µg/ml normocin, 20ng/mL FGF, 20ng/mL EGF, and 20ng/mL of PDGF-AA. Mouse mIDH1 NS were grown in DMEM-F12 media supplemented with 100units/mL antibiotic-antimycotic, B27, N2, 100µg/mL normocin, 10ng/mL FGF and 10ng/mL EGF. Stable generation of ZMYND8 KO glioma cells were maintained in culture with 10ug/mL puromycin. Glioma cells were dissociated using StemPro Accutase solution and passaged every 4 days for mouse NS and weekly for human glioma cells. Human glioma cells were shared through the following collaborations: Dr. Daniel Cahill laboratory, Harvard Medical School (MGG119) and Dr. Joseph Costello laboratory, UCSF (SF10602). LC1035 were generated in our laboratory through our collaboration with the University of Michigan’s Department of Neurosurgery.

### RNA-sequencing (RNA-seq)

Bulk RNA-Sequencing was performed in collaboration with the University of Michigan (UM) Advanced Genomics Core (AGC), using the HiSeq-4000 platform for 150-base paired-end reads. Total RNA was isolated from patient-derived mIDH1 GCC SF10602 untreated and AGI-5198- treated using the RNeasy Plus Mini Kit and 100ng of RNA was submitted to the UM AGC. RNA quality was assessed using the TapeStation, samples with an RIN (RNA Integrity Number) of 8 or greater were prepared using Illumina TruSeq mRNA Sample Prep Kit v2. cDNA Library preparation was performed by UM DSC using a ribo-depleted RNA protocol method. Results are from 3 technical replicates per condition; SF10602 untreated or treated with mIDH1 inhibitor AGI- 5198 for 1 week. Information pertaining to the number of aligned reads utilized for our RNA-seq analysis can be found in Supplementary Table 1.

### Bioinformatics analysis of RNA-seq for differential expression

Illumina HiSeq read files were downloaded to the Sequencing Core storage and concatenated into a single .fastq file for each sample. In collaboration with the UM Bioinformatics Core, we checked the quality of the raw reads data for each sample using FastQC (version v0.11.3) to identify features within the data that may indicate quality issues (e.g. low quality scores, over-represented sequences, inappropriate GC content). We aligned reads to the reference genome (UCSC hg19) using Bowtie2 (version 2.2.1). Expression quantitation was performed with HTSeq (version 0.6.1), set to count non-ambiguously mapped reads only. Data were pre-filtered to remove genes with 0 counts in all samples. Normalization and differential expression was performed with DESeq2 (version 1.14.1), which uses a negative binomial generalized linear model. Resultant p-values were corrected for multiple hypotheses using the Benjamini-Hochberg False Discovery Rate (FDR). Genes and transcripts were identified as differentially expressed based on FDR ≤ 0.05, and fold change ≥ ± 1.5. The volcano plot was produced using an R base package and encompasses all genes identified by our RNA-seq analysis.

### GSEA Enrichment Map and Gene Ontology (GO) Analysis

All 1335 differentially expressed genes between SF10602 treated with AGI-5198 (mIDH1 specific inhibitor) vs. untreated SF10602 were exported to a table with their gene names in the first column and log2-fold change values in the second column, sorted to produce gene ranks. The rank file was used as input to the GSEA (Gene Set Enrichment Analysis), downloaded from the Broad Institute (http://www.gsea-msigdb.org/gsea/login.jsp). The Broad Institute MSigDB includes the complete Gene Ontology (GO) (c5.all.v7.5.1.symbols.gmt), which we used as the input Knowledgebase. GSEA was run with 2000 permutations and gene set size range restricted to between 0 and 200 genes. An enrichment map was generated using the Cytoscape Platform (v3.9.1), which requires the rank and gmt file along with the positive/negative GSEA results. Each circle represents an individual GO term, and the size of the circle represents the number of genes within that set. The color of the circle signifies the directionality of expression in the mIDH1 inhibitor treated (AGI- 5198) compared to untreated, where red is upregulated in AGI-5198-treated compared to untreated while blue is downregulated. We set a stringent cut-off with a log2FC >±0.6, p-value<0.003, and FDR<0.002 shown in Figure S1A. The enrichment score plots for Replication Fork, DNA Replication Initiation and Nuclear Replication Fork are shown in Figure 1E, which are images provides in the GSEA Pre-ranking report.

**Figure 1:**
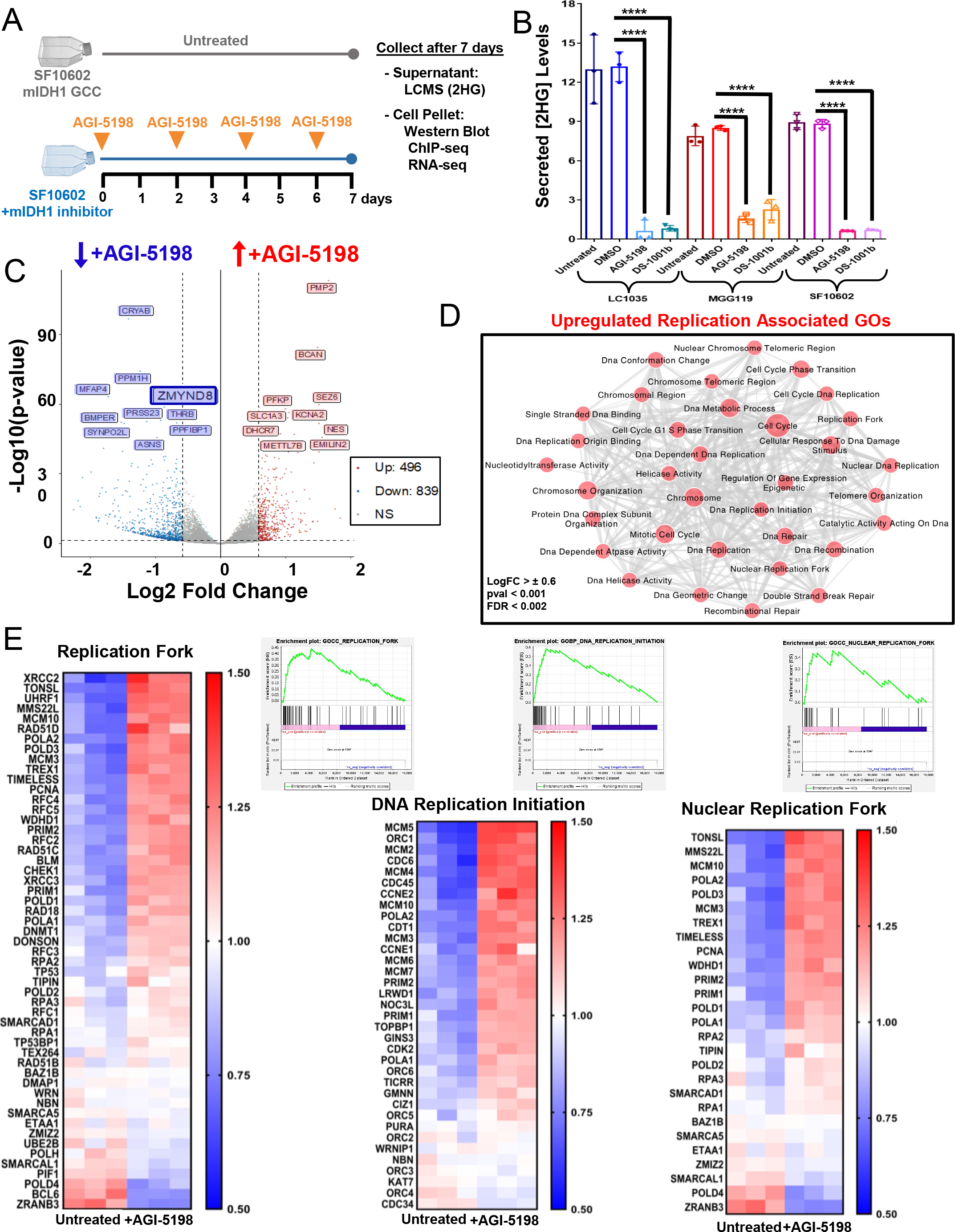
Inhibition of mIDH1-R132H in a patient-derived glioma cell culture (SF10602) leads to increased activation of DNA damage associated gene ontologies (GO). (**A**) Experimental schematic of downstream analyses comparing untreated SF10602 human glioma cell culture (GCC) that endogenously express the R132H mutation in isocitrate dehydrogenase 1 (mIDH1) with cells treated with 5μM AGI-5198 (a mIDH1 specific inhibitor). After 1 week, the supernatant was collected for liquid chromatogram and mass spectrometry (LCMS) and cell pellet was utilized for transcriptomic (RNA-sequencing), epigenomic (ChIP- sequencing) and protein (Western blot) analysis. (**B**) Representative histogram displaying the quantification of 2-hydroxyglutarate (2HG) using LCMS, an oncometabilite produced by mIDH1, present within the conditioned media of three patient-derived mIDH1 GCCs [LC1035, MGG119 and SF10602]. Samples were collected from untreated, vehicle treated (DMSO), mIDH1 inhibitor (AGI-5198) or clinical mIDH1 inhibitor (DS-1001b). Errors bars represent standard error of mean (SEM) from independent biological replicates (n=3). **** P < 0.0001; unpaired t test. (**C**) Volcano plot of differentially expressed genes (DEG) based on RNA-seq analysis comparing AGI-5198 treated SF10602 vs. untreated. Dots represent individual genes, genes found to be downregulated in AGI-5198 treated mIDH1 GCCs vs. untreated are shown in blue, up-regulated in red, or no statistically significant (NS) difference in grey. (**D**) Gene Set Enrichment Analysis (GSEA) map of upregulated GOs in AGI-5198 treated GCCs were associated with replication. (**E**) Heatmap depicting the relevant GOs related to DNA replication initiation and replication forks that were upregulated in AGI-5198 treated GCCs compared to untreated. The corresponding GSEA enrichment plot is adjacent, where the green line represents the enrichment score for a given GO as the analysis proceeds along the rank list of DEG.

### ChIP-seq

SF10602 were cultured in the presence of 5µM AGI-5198 (mIDH1 inhibitor) or the vehicle (DMSO) for 10 days prior to isolation for chromatin immunoprecipitation (ChIP). Modifications to laboratory native ChIP-seq protocol as previously described in Mendez et al. JoVE 2018 (Nunez et al. 2019) were as follows: 1.) increased cell number per histone mark IP from 1x10^6^ to 3x10^6^, 2.) increased histone mark antibody concentration to 2µg, 3.) extended Dynabead A/G incubation to 6hrs, and 4.) doubled Dynabead A/G concentration.

### Lentiviral particle generation

Second generation lentiviral particles were packaged using HEK293T cells, which were seeded at 5 million cells and incubated with plasmid transfection mix containing 80µl of jetPRIME solution, 2 mL of PEG, envelope (10µg), package (15µg) and lentiCRISPR-V2-ZMYND8 (20µg) plasmid provided by Dr. Weibo Lou at UTSW. (Wang et al. 2021 Cancer Research) Lentiviral particles were purified from conditioned media by centrifuging at 30,000 rpm for 2hrs. After ultra- centrifugation, the concentrated lentiviral particles were resuspended in 1 mL of ice-cold PBS under gentle agitation 25rpm at 4°C for 1hr. Lentiviral particle solution was aliquoted into cryo-safe microcentrifuge tubes and stored at -80°C for later use.

### Intracranial mIDH1 glioma model

Our laboratory has modelled mIDH1 low grade glioma by integrating oncogenic plasmid DNA into the neural stem cells present within the developing brain of post-natal mice utilizing the sleeping beauty (SB) transposon system. The mIDH1 glioma cells endogenous express IDH1- R132H along with genetic lesions that drive oncogenesis through NRAS^G12V^ and simulate tumor suppressor loss by ATRX and TP53 short hairpin knockdown. Tumor NS were derived from endogenous mIDH1 tumors that were adapted to *in vitro* culture and could be reimplanted into immunocompetent C57BL6 mice for preclinical experiments in this study. Intracranial surgeries were performed by stereotactically injecting 50,000 mIDH1 NS into the right striatum using a 22- gauge Hamilton syringe with the following coordinates: 1.0 mm anterior, 2.5mm lateral and 3.0mm deep from the brega suture line.

### Generation of human and mouse ZMYND8 KO glioma cells

Human mIDH1 primary glioma cell cultures SF10602 and MGG119 were seeded at 2x10^5^ cells per well of laminin coated 6-well plate. The following day, cell culture media was removed and cells were incubated directly with lentiviral particles for 10mins. Media was changed after 3 days of lentiviral transfection and cells were expanded for another week prior to puromycin selection. Loss of ZMYND8 expression was confirmed by WB.

### Embedding mIDH1 GCC NS for Immunohistochemistry

Following 1 week treatment with vehicle (DMSO) or 5μM mIDH1 inhibitor (AGI-5198), mIDH1 NS were transferred to 15 mL conical and allowed to settle by gravity. The supernatant was removed by manual pipetting and mIDH1 NS were suspended gently in 4% PFA for overnight incubation to fix and retain neurosphere morphology. The next day mIDH1 NS were washed thrice in PBS buffer. Autoclaved 4% low-melting agarose was cooled to 43°F on bead bath. In order to embed mIDH1 NS in agarose, cells were transferred to 1.5mL microcentrifuge tube and 200µl of agarose was added over top of the NS. A wooden skewer was used to evenly distribute mIDH1 NS before immediately transferring to ice. Agarose cone containing mIDH1 NS was removed and processed at Histology core for paraffin sectioning. Representative sections of mIDH1 NS were made at 5μm thickness and transferred to microscope slides. Immunohistochemistry (IHC) was performed by heating slides to 60°C for 20mins to assist in remove of paraffin. NS slides were deparaffinized and rehydrated. Permeabilization of NS was performed using TBS-0.025% Triton-X (TBS-TX) for 20 min. Antigen retrieval was performed at 96°C with citrate buffer (10mM sodium citrate, 0.05% Tween-20, pH 6) for an additional 20 min. Once cooled to room temperature (RT), sections were outlined with hydrophobic barrier pen, washed thrice (3min washes) with TBS-TX and blocked with 10% horse serum (HS) for 1hr at RT. IHC sections were incubated in primary antibody ZMYND8 (1:2000) diluted in 5% HS TBS-TX overnight in 4°C cold room. The following day, sections were washed with TBS and incubated with biotinylated secondary antibody goat anti-rabbit (1:1000) for 1hr at RT. Next, slides were washed thrice in TBS and incubated with Vectastain ABC reagent for 30mins covered by aluminum foil. Following TBS wash, slides were developed using Betazoid DAB Chromogen kit (Biocare BDB2004) for 1-2mins at RT.

### *In vitro* Dose-Response and evaluation of radiosensitivity using PARP inhibitor

To assess the susceptibility of both ZMYND8 WT and ZMYND8 KO generated in mouse mIDH1 NS and human mIDH1GCCs to PARP inhibitor (PARPi), we used Pamiparib (BGB-290) (SelleckChem, S8592), a PARPi for our study. Herein, both the cells were plated at a density of 1500 cells per well in 96-well plates (Fisher, 12-566-00) 24h prior to treatment. We used 5 wells per inhibitor dose evaluated for each cell type. We evaluated 8 concentrations of inhibitor in serial dilutions (e.g., 1µM, 3µM, 10µM, 30µM, 50µM, 100µM) and were added to each well for each dilution evaluated. Cells were then incubated for 3 days and viability was assessed using CellTiter-Glo assay (Promega, G7572) following manufacturer’s protocol. Moveover to assess the radiosensitization, cells were then incubated with either Pamiparib alone or in combination with radiation at their respective IC_50_ doses for 72h in triplicate wells per condition. Cells were pre-treated with PARPi 2h prior to irradiation with 6Gy and 20Gy of radiation for mouse neurospheres and human glioma cells respectively. Resulting luminescence was read with the Enspire Multimodal Plate Reader (Perkin Elmer). Data was represented graphically using the GraphPad Prism 7 using sigmoidal regression model which allows the determination of IC_50_ values and statistical significance was determined by one-way ANOVA followed by Tukey’s test for multiple comparisons.

### Western blot and radiation sensitivity *in vitro*

Mouse mIDH1 NS (NPAI) and human mIDH1 GCCs (SF10602 and MGG119) were seeded at density of 5.0 x 10^6^ cells into 75-cm2 flasks containing NSC media. After 24 hours, mouse NS and human GCCs were treated with 5Gy and 20Gy radiation respectively. After an additional 48 hours, cell lysates were prepared by incubating glioma cells with RIPA lysis buffer (MilliporeSigma, R0278) and 1X Halt™ protease and Phosphatase inhibitor cocktail, EDTA-free (100X) (Thermo Scientific, 78441) on ice for 15 minutes. Resulting cell lysates were centrifuged at 14000 RPM at 4°C for 15 minutes and supernatants were collected to determine protein concentration in comparison to standard bovine serum albumin (BSA) protein concentrations through bicinchoninic acid assay (BCA) (Pierce, 23227). For electrophoretic separation of proteins, 20 μg of total protein were resuspended in loading buffer (10% sodium dodecyl sulfate, 20% glycerol, and 0.1% bromophenol blue) and samples were heated at 95°C for 5mins to denature protein and later loaded onto a 4-12% Bis-Tris gel (Thermo Fisher Scientific, NuPAGE, NP0322BOX). Proteins from the gel were transferred to 0.2 μm nitrocellulose membrane (Bio-Rad, 1620112) and blocked with 5% bovine serum albumin (BSA) in TBS-0.1% Tween-20. After blocking, membranes were incubated with primary anti-phospho γH2AX (1:1000) (Cell Signaling Technologies, 9718S), anti-γH2AX (1:1000) (Cell Signaling Technologies, 2595S), Vinculin (1:3000) (Fisher Scientific, PI700062) or β-actin antibodies (1:4000) (Sigma-Aldrich, A1978) overnight at 4°C. The next day, blots were washed with TBS-0.1% tween-20 and incubated with secondary (1:4000) antibodies [Dako, Agilent Technologies, goat anti-rabbit 1:4000 (P0448), rabbit anti-mouse 1:4000 (P0260)] for one hour at room temperature. Blots were washed several times again with TBS-0.1% tween-2.0 Enhanced chemiluminescence reagents were used to detect the signals following the manufacturer’s instructions (SuperSignal West Femto, Thermo Fisher Scientific, 34095) and visualized under Bio-Rad gel imaging software. Band intensities were quantified using ImageJ software. The same procedure was performed for Chk1, pChk1, Rad51, pRad51, ATM, pATM, FANCD2 and pFANCD2 proteins using the antibodies and dilutions described in table SX.

## Results

### Inhibition of mIDH1 by AGI-5198 leads to the upregulation of replication associated pathways

In this study, we sought to determine radioresistance mechanisms in IDH-mutant gliomas that contribute to glioma cell survival in response to radiotherapy. We utilized a radioresistant patient-derived glioma cell culture SF10602, which retained the endogenous mutations in IDH1- R132H, ATRX and TP53 after adaptation to *in vitro* culture. (27) In order to identify pathways altered by mIDH1 inhibition in SF10602, we performed a transcriptomic (RNA-seq) and epigenomic (ChIP-seq) screen (Figure 1A) by administering 5µM AGI-5198 every 2 days for 1 week. It has been shown that 2HG is released within the conditioned media of mIDH1 GCC maintained *in vitro*. (28) To confirm that our mIDH1 inhibitor treatment schedule inhibited 2HG production, we collected the conditioned media from untreated and AGI-5198 treated mIDH1 GCCs to analyze the presence of 2HG by liquid chromatogram and mass-spectrometry (LCMS). We observed a significant reduction of 2HG levels (Figure 1B) in three mIDH1 GCCs SF10602, LC1035 and MGG119 following treatment with AGI-5198 or another mIDH1 inhibitor DS-1001b currently in phase II clinical trials (NCT04458272) as compared to the vehicle control DMSO.

Inhibition of mIDH1 by AGI-5198 resulted in dynamic changes in transcriptional regulation (Figure 1C), where we observed 1,335 differentially expressed genes consisting of 498 upregulated and 839 downregulated genes based on a log2 fold-change (log_2_FC) cut-off greater than **±**0.6. We performed a gene set enrichment analysis (GSEA) of these differentially genes altered by mIDH1 inhibition to uncover pathways dependent upon mIDH1 reprogramming (Supplemental Fig. S1A). We observe an upregulation of gene ontologies (GO) associated with replication (Figure 1D) specifically Replication Fork (GO:0005657), DNA Replication Initiation (GO:0006270) and Nuclear Replication Fork (GO:0043596). Many of the genes shared across these replication associated GOs have roles in regulating replication stress facilitated by the stalling and restart of replication forks (Figure 1E). In addition, the genes that were upregulated following mIDH1 inhibitor treatment function as part of the core replisome or have been associated with responding to replicative stress. This data suggests that blocking 2HG production in mIDH1 GCCs using AGI- 5198 induces replicative stress.

### Epigenetic Changes following mIDH1 inhibition suggests activation of replication stress

When cells divide, there must be a strict coordination between transcription and replication to ensure the maintenance of genomic stability. This tightly regulated process is disrupted in cancer cells as a result of oncogene activation. (29) The destabilization of replication forks or collision events between mediators of transcription and replication can generate DSB. To address replication stress, cells express proteins involved in cell cycle arrest, replication fork restart, and DNA repair to resolve damaged DNA regions. (29) In order to translate the elevated expression of replication stress genes to their regulation at the chromatin level, we performed chromatin immunoprecipitation and deep sequencing (ChIP-seq) for histone marks associated with active transcription (H3K4me3, H3K36me3), enhancers (H3K4me1, H3K27ac) and transcriptional repression (H3K27me3) in SF10602 in the presence of AGI-5198 (Figure 2A).

**Figure 2:**
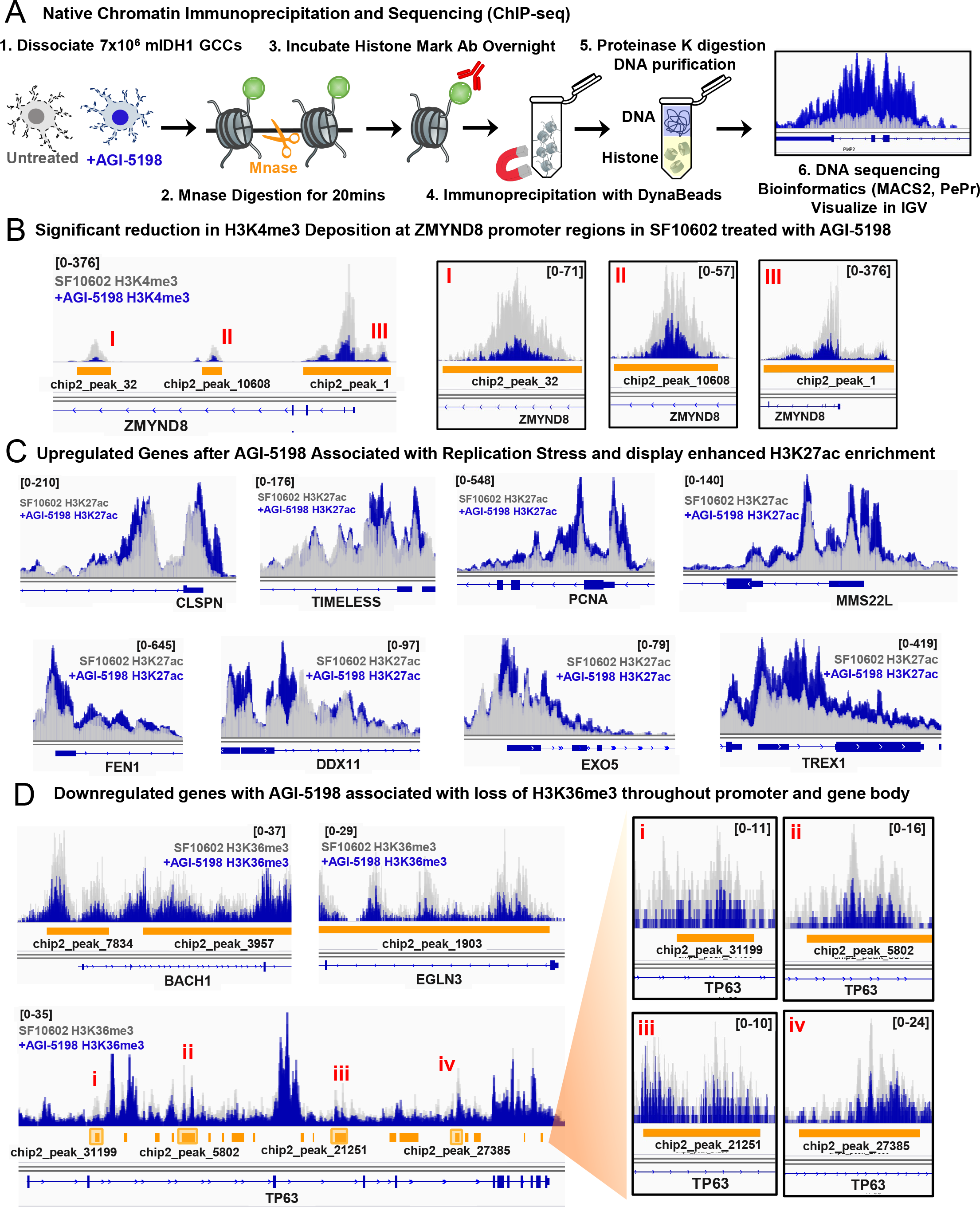
Epigenomic changes in histone mark deposition following mIDH1 inhibition. (**A**). Diagram detailing the steps of native chromatin immunoprecipitation and sequencing (ChIP-seq) (**B**) Integrative Genomics Viewer (IGV) image displays overlapping tracks comparing the H3K4me3 occupancy in specific genomic regions near the ZMYND8 promoter in untreated SF10602 (grey, two replicates) and AGI-5198 treated (dark blue, two replicates). The y-axis represents the number of immunoprecipitated fragments for a given histone mark normalized to the total number of reads per sample and mapped to the human genome reference (hg19) along the x-axis. Regions that display significant changes in histone mark deposition based on peak-calling prioritization pipeline (PePr) comparison between SF10602 untreated vs. AGI-5198 treated SF10602 are represented by orange bars and red roman numerals signify distinct regions that are expanded to the right. (**C**) IGV screenshots of H3K27ac deposition at the promoters of specific genes associated with replication stress. (**D**) IGV screenshots of H3K36me3 deposition throughout the promoter and gene body, where select regions that lost H3K36me3 deposition by PePr calling at TP63 locus are indicated by yellow squares.

We focused on mediators of the HR pathway, we found ZMYND8 to one of the most significantly downregulated genes following the treatment of SF10602 with AGI-5198 (Figure 1C). The chromatin reader ZMYND8 has been shown to be recruited to DSB at actively transcribed regions. (12) Depletion of ZMYND8 by siRNA or knockout impaired the recruitment of RAD51 foci at sites of DNA damage. (12, 15) We observed a significant loss of the active histone mark H3K4me3 at three distinct regions (I, II, III) near the promoter of ZMYND8 following mIDH1 inhibition by AGI-5198 (Figure 2B).

To determine if the upregulation of ZMYND8 was dependent on mIDH1 expression, we assessed nascent transcription by bromouridine sequencing (Bru-seq) within our genetically engineered mouse glioma model comparing mouse tumor NS clones expressing either wtIDH1 (NPA) or mIDH1 (NPAI) (Supplemental Figure 2A). Mouse mIDH1 NS displayed an enhanced transcription rate for *ZMYND8* (1.27-fold), along with HR proteins mediators *RAD50* (1.78-fol*d), FANCA* (1.63-fold) and *RAD51*(1.30-fold) compared to the wtIDH1 NS. (Supplemental Figure 2B). To determine whether there were differences in the epigenetic regulation of ZMYND8 in the mIDH1 NS compared to wtIDH1 NS, we analyzed H3K4me3 enrichment at the ZMYND8 locus. We found five PePr differential enriched peaks (i-v) that showed significantly higher H3K4me3 enrichment corresponding to ZMYND8 promoter regions across the mIDH1 NS (NPAI C1, C2, C3) versus wtIDH1 NS (NPA C54A, C54B, C2) clones (Supplemental Figure 2C). These findings suggest that mIDH1 epigenetically regulates ZMYND8 by increasing H3K4me3 deposition at the promoter leading to enhanced transcription.

Next, we investigated the epigenetic changes that occurred at genes associated with replication stress. We observed elevated H3K27ac deposition at the promoter regions of genes associated with replication following AGI-5198 treatment (Figure 2C). The progression of replication forks is facilitated by the unwinding of DNA by the replisome, which is comprised of claspin (CLSPN) and Timeless (TIMELESS) and the tethering of DNA polymerase by proliferating cell nuclear antigen (PCNA). (30) Methyl methanesulfonate-sensitivity protein 22- like (MMS22L) functions in a complex with Tonsoku Like (TONSL), where it has been shown to be required for replication fork stability by binding newly incorporated histones and aiding in RAD51 loading. (31) The flap endonuclease 1 (FEN1) resolves nicks in DNA that arise during replication. (32) The DEAD/H-BOX Helicase 11 (DDX11) facilitates the progression of replication forks by unwinding G-quadraplexes. (33) The endonuclease V (EXO5) supports replication fork restart and EXO5 expression is correlated with higher mutational burden in solid tumors. (34) Three Prime Repair Exonuclease 1 (TREX1) degrades cytosolic DNA released from the nucleus as a result of chromosomally unstable cells. (35) These data suggest that mIDH1 inhibition enhances the activation of replication stress genes marked by elevated deposition of regulatory histone mark H3K27ac at the promoter regions.

When we examined the chromatin regions of genes downregulated with AGI-5198 treatment and were associated with replication, we observed loss of H3K36me3 deposition throughout the promoter and gene body. (Figure 2D) BTB Domain and CNC Homolog 1 (BACH1) has a role in activating ATR-dependent Chk1 phosphorylation. (36) Egl-9 Family Hypoxia Inducible Factor 3 (EGLN3) drives ATR-mediated repair under hypoxic conditions. (37) TP63 regulates transcription at tissue-specific enhancers and conditional ablation of p63 lead to apoptosis of adult neural precursors cells in postnatal mice. (38) These data suggest transcriptional suppression of proteins involved in ATR activation following mIDH1 inhibition.

### Inhibiting mIDH1 suppresses protein expression of ZMYND8

Epigenetic regulation of a gene is defined by the patterns of specific histone modifications at the genomic locus and correlates with chromatin accessibility and gene expression. H3K4me3 is enriched at sites of poised and actively transcribed regions, which highlights the activity of genes that govern cell-specific networks. We sought to determine if the significant reduction of H3K4me3 at the ZMYND8 promoter following AGI-5198 treatment in the SF10602 would lead to a decrease in protein expression. We included two additional human mIDH1 GCCs (MGG119, LC1035) to evaluate if the inhibition of mIDH1 lead to a decrease in ZMYND8 gene expression across mIDH1 GCCs derived from separate patients. We performed western blotting (WB) for ZMYND8 in three human mIDH1 GCCs treated with AGI-5198 (Figure 3A). We observed a significant reduction of ZMYND8 protein expression in the MGG119 (*P<0.001*), SF10602 (*P<0.0001*) and LC1035 (*P<0.001*) following AGI-5198 treatment (Figure 3B). To confirm that this modulation of ZMYND8 protein expression was consistent with mIDH1 inhibition, we treated the human mIDH1 GCCs with DS-1001b (Figure 3C). We observed a significant reduction of ZMYND8 expression in the MGG119 (*P < 0.05*), SF10602 (*P < 0.0001*) and LC1035 (*P<0.01*) after DS-1001b treatment (Figure 3D). Considering that western blots are representative of the bulk sample, we performed immunohistochemistry for ZMYND8 on paraffin-embedded human mIDH1 NS pre-treated with AGI-5198. This allowed us to determine the impact of mIDH1 inhibition on ZMYND8 expression at the individual cell level (Figure 3E). We analyzed the nuclear IHC staining of ZMYND8 using Quantitative Pathology & Bioimage Analysis (QuPath) to define the number of ZMYND8 positive cells per frame (Supplemental Figure 3A). We observed a significant reduction in the percentage of ZMYND8 positive nuclei in the SF10602 (*P< 0.001*) and LC1035 (*P < 0.0001*) pre-treated with AGI-5198 compared the DMSO treated (Figure 2F). Pre-treatment of LC1035 with DS-1001b prompted a similar reduction in the percentage of ZMYND8 positive nuclei (Supplemental Figure 3B). These data demonstrate that mIDH1 epigenetically regulates ZMYND8 protein expression, which is suppressed by mIDH1 inhibition.

**Figure 3:**
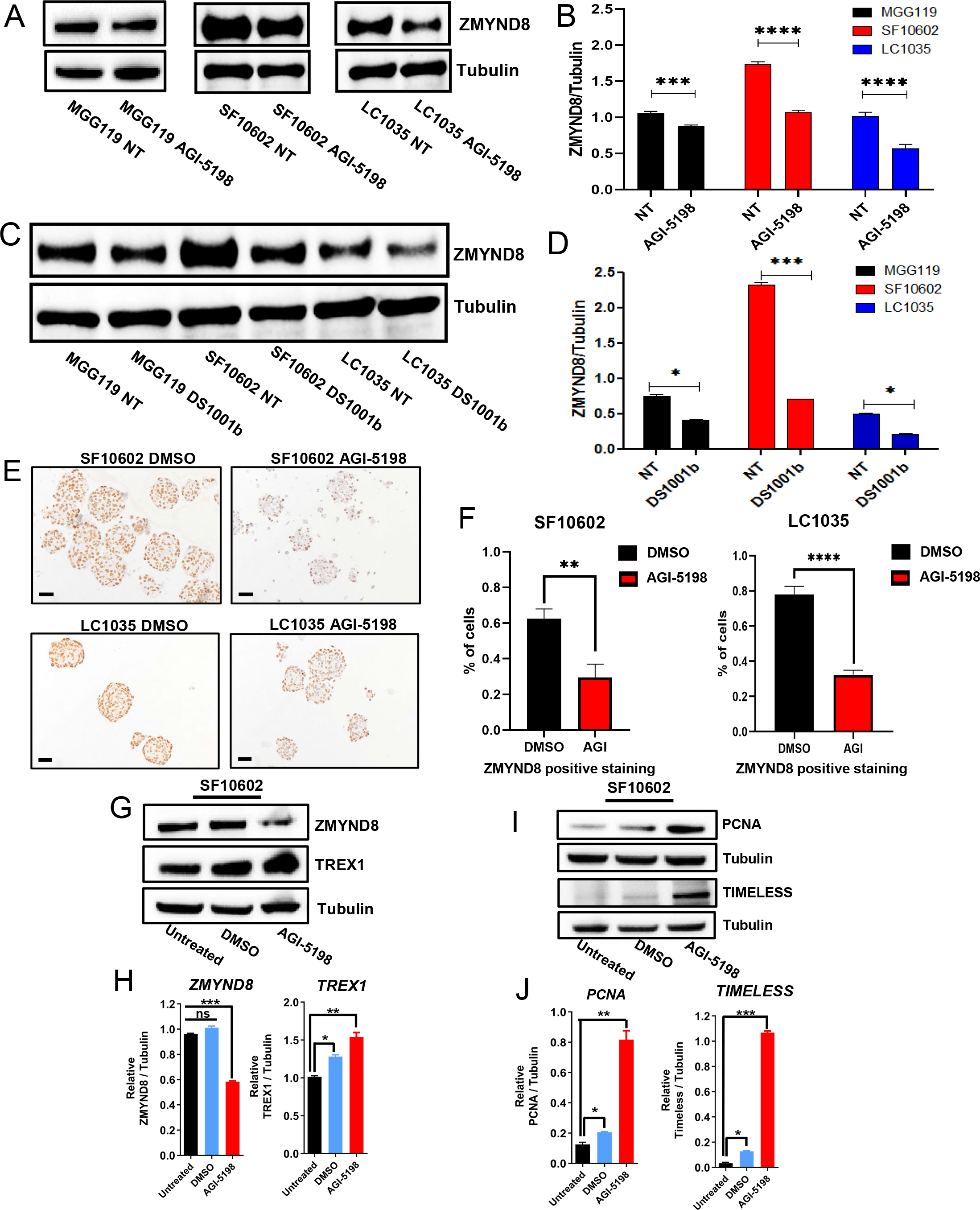
Decreased ZMYND8 protein expression following mIDH1 inhibitor treatment (AGI-5198, DS1001b) in mIDH1 GCCs, coincides with an increased expression of genes associated with genomic instability. (**A**). Representative western blot for ZMYND8 protein expression in three human mIDH1 GCCs that were non-treated (NT) or AGI-5198-treated (mIDH1 inhibitor) for 1 week. (**B**) ImageJ densiometric quantification of ZMYND8 protein expression based relative to loading control (tubulin) for each of the mIDH1 GCCs either NT or AGI-5198 treated: MGG119 (black), SF10602 (red), and LC1035 (blue). (**C**) Representative western blot for ZMYND8 protein expression in three human mIDH1 GCCs that were NT or DS1001b treated (clinical mIDH1 inhibitor) for 1 week. (**D**) ImageJ densiometric quantification of ZMYND8 protein expression following DS1001b treatment compared to NT. (**E**) Immunohistochemistry (IHC) staining of sectioned paraffin-embedded human mIDH1 GCCs treated with vehicle (DMSO) or mIDH1 inhibitor (AGI-5198). (**F**) Representative Quantitative Pathology & Bioimage Analysis (QuPath) of IHC slides to identify the percentage of ZMYND8 positive staining (Diaminobenzidine (DAB) optical density > 0.4) for 12 representative frames. (**G**) Western blot analysis shows ZMYND8 and TREX1 expression in SF10602 mIDH1 GCC either untreated, DMSO, or AGI- 5198 treatment for 1 week with tubulin as a loading control. (**H**) ImageJ densiometric quantification of the western blot for ZMYND8 and TREX1. (**I**) Western blot analysis shows PCNA and TIMELESS expression in SF10602 either untreated, DMSO, or AGI-5198 treatment for 1 week with tubulin as a loading control. (**J**) ImageJ densiometric quantification of the western blot for PCNA and TIMELESS. Errors bars represent SEM from independent biological replicates (n=3). * P<0.05, ** P<0.01, *** P<0.001, **** P < 0.0001; unpaired t test

The reproducible reduction of ZMYND8 protein expression following mIDH1 inhibition led us to explore whether the decrease in ZMYND8 coincided with changes in proteins involved in replication stress (Figure 3G). To address this, we collected protein from SF10602 that were untreated, treated with vehicle (DMSO) or 5µM AGI-5198 every 2 days for 1 week. First, we investigated the protein expression of TREX1, an exonuclease that sequesters ssDNA fragments generated from aberrant replication and has been shown to be recruited to stalled replication forks (Figure 3G). (39) We observed a significant increase in TREX1 protein expression in SF10602 treated with AGI-5198 (*P > 0.01*) compared to untreated cells, and a slight increase (*P > 0.05*) in DMSO-treated cells (Figure 3H). Next, we evaluated the expression of PCNA and TIMELESS, which are known to be co-expressed during S phase (Figure 3I). (40) We observed a significant increase in PCNA protein expression in SF10602 treated with AGI-5198 (*P > 0.01*) compared to untreated cells, and a modest increase (*P > 0.05*) in DMSO-treated cells (Figure 3H). Similarly, we found TIMELESS to be significantly increased following AGI-5198 treatment (*P > 0.001)* compared to untreated SF10602. In order to maintain the structural stability of stalled replication forks, Timeless forms a complex with Tipin to prevent the disassembly of the replisome.(40) These findings suggest that mIDH1 inhibition induces genomic instability mediated by an increase in replication stress protein expression.

### Loss of ZMYND8 enhances the radiosensitivity of mIDH1 GCCs to irradiation and exploits the vulnerability of mIDH1 GCCs to epigenetic inhibitors targeting BRD4 and HDAC

Based on the role of ZMYND8 in transcriptional repression at sites of DSB, we hypothesized that it might promote resistance to IR-induced DNA damage in mIDH1 GCCs. We began by targeting the human and mouse isoforms of ZMYND8 using short-hairpin (shRNA) knockdown that we cloned into our pT2-plasmid backbone (Supplemental Figure 4A-B). (41, 42)To confirm that the pT2-shZMYND8-GFP plasmids we designed could reduce ZMYND8 protein expression, we transfected HEK293 cells with pT2-shZMYND8-GFP plasmids targeting the human gene and NIH-3T3 cells for the mouse isoform in combination with the sleeping beauty transposase (SB) plasmid to improve the integration efficiency. (9) We performed WB analysis 3 days post-transfection and observed a reduction in ZMYND8 expression in HEK293 transfected with human sh2-ZMYND8 and mouse sh4-ZMYND8 plasmids in NIH3T3 (Supplemental Figure 4C).

Next, we generated stable ZMYND8 shRNA knockdown clones in two mIDH1 GCCs, SF10602 and MGG119. (Supplemental Figure 5A). After purifying the GFP+ population by flow cytometry, we assessed ZMYND8 protein expression by WB in the MGG119 and SF10602 (Supplemental Figure 5B and 5E). We observed a significant reduction of ZMYND8 protein expression by 75% in the MGG119 and 60% in the SF10602 compared to non-transfected (Supplemental Figure 5C and 5F). Survival of mIDH1 GCCs expressing the shZMYND8-GFP plasmids was assessed 3 days post-irradiation exposure. MGG119 expressing the shZMYND8- GFP plasmids showed reduced cellular viability in response to escalating doses of radiation with a half maximal inhibitory concentration (IC_50_) of 9.5Gy (Supplemental Figure 5D). SF10602 expressing shZMYND8-GFP plasmids displayed a significant reduction in cellular viability but a IC_50_ was not reached at single radiation doses up to 20Gy (Supplemental Figure 5G). These data suggest that suppression of ZMYND8 expression in mIDH1 GCC enhances their susceptibility to ionizing radiation.

Next, we ablated the expression of ZMYND8 using lentiviral CRISPRv2 vector expressing Cas9 and ZMYND8 guide RNAs (sgRNA) to generate ZMYND8 KO mIDH1 GCC clones (Figure 4A). (20) After lentiviral incubation and puromycin selection, loss of ZMYND8 was evaluated by WB in the SF10602, MGG119 and NPAI mouse mIDH1 NS (Figure 4B, 4D, and 4F respectively). We observed a significant reduction in cellular viability 3 days post-IR in the ZMYND8 KO compared to ZMYND8 WT mIDH1 GCCs. The SF10602 ZMYND8 KO had an IC_50_ of 7.5Gy compared to SF10602 ZMYND8 WT where an IC_50_ was not reached (Figure 4C). The MGG119 ZMYND8 KO had an IC_50_ of 16.8Gy compared to MGG119 ZMYND8 WT where an IC_50_ was not reached (Figure 4E). The NPAI ZMYND8 KO exhibited an IC_50_ of 2.8Gy versus the NPAI ZMYND8 WT where an IC_50_ was not reached. (Figure 4G). These data indicate ZMYND8 contributes to the survival of mIDH1 GCCs in response to radiation.

**Figure 4:**
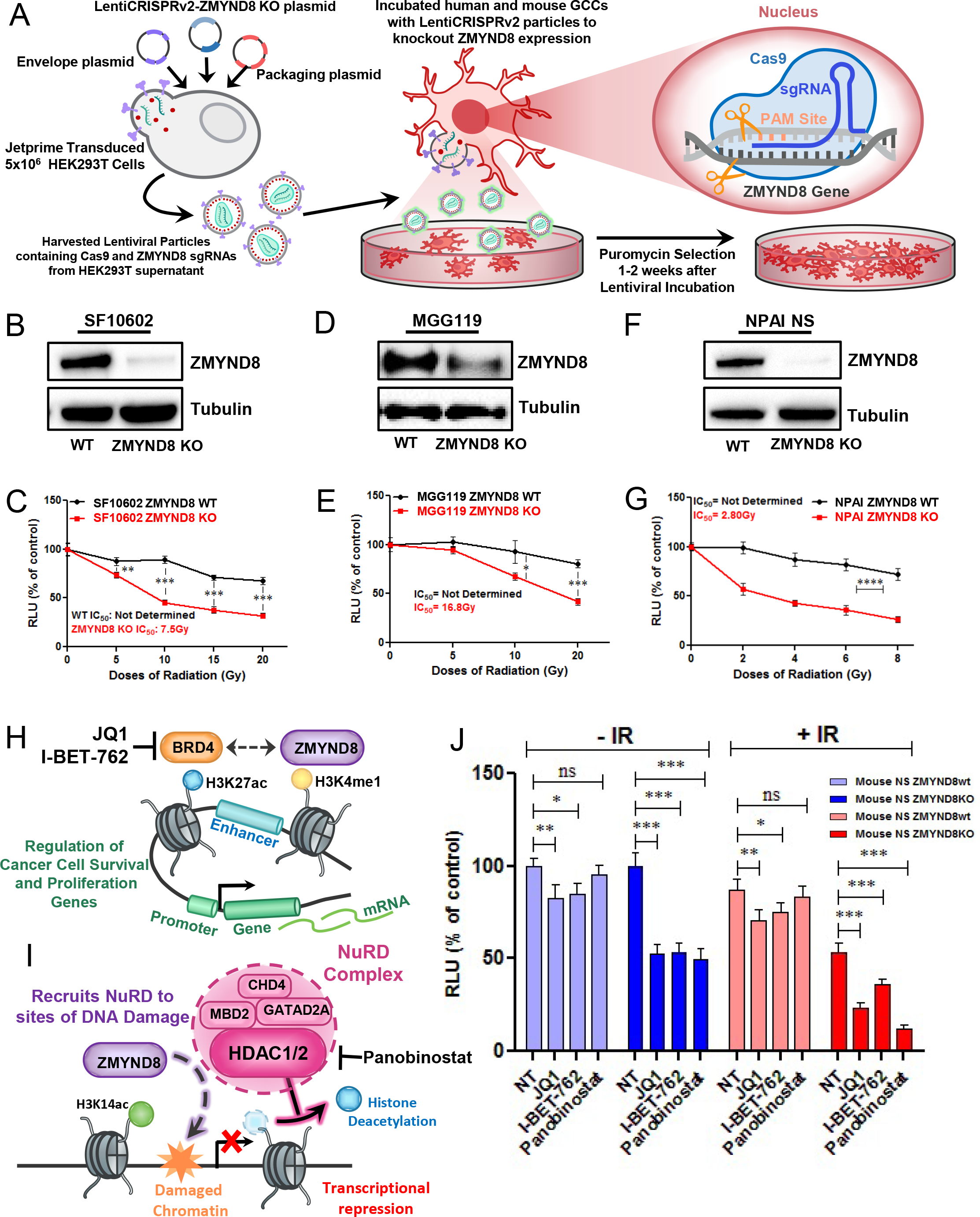
ZMYND8 Knockout GCCs display reduced viability to irradiation, which is further enhanced when combined with BRD4 and HDAC inhibition. (**A**). Experimental model in which ZMYND8 lentiviral particles were generated to knockout (KO) ZMYND8 expression in human and mouse GCCs mediated by CRISPR-Cas9-sgRNAs. ZMYND8 KO GCCs were selected based on resistance to 10μg/mL puromycin for 1 week in mouse mIDH1 GCCs and 2 weeks for human mIDH1 GCCs. Representative western blot quantification of ZMYND8 KO for (**B**) SF10602, (**D**) MGG119 and (**F**) NPAI mouse NS. Cellular viability of ZMYND8 wildtype (WT, shown by black line) vs. ZMYND8 KO, represented by the red line, was assessed 72 hours after irradiation (IR) exposure using CellTiter-Glo assay in the (**C**) SF10602, (**E**) MGG119 and (**G**) NPAI. Results are expressed in relative luminescence units (RLU) to control non-irradiated (0 Gy) cells. (**H**) Working model of ZMYND8-interacting partner Bromodomain-containing protein (BRD4), where ZMYND8 is recruited to enhancer regions marked by H3K4me1 and contributes to the regulation of cancer cell survival and proliferation associated genes. BRD4 inhibitors, JQ1and I-BET-762, can disrupt this interaction. (**I**) Working model of ZMYND8-interacting partner HDAC1/2 (histone deacetylase 1/2); a component of the Nucleosome Remodeling and Histone Deacetylase (NuRD) complex , where ZMYND8 binds H3K14ac residues present at damaged chromatin regions and recruits HDAC along with other NuRD subunits:MDB2, CHD4, and GATAD2A. Panobinostat inhibits HDAC1/2 and prevents histone deacetylation medidated by HDAC1/2, which is required for transcriptional repression at regions of DNA damage. (**J**) Representative bar graph of cellular viability measured in RLU, which shows the effect of BRD4 (JQ1, I-BET-762) or HDAC (Panobinostat) inhibition alone (-IR) or in combination with irradiation (+IR) in mouse NPAI ZMYND8 wt in blue vs. NPAI ZMYND8 KO in red. Errors bars represent SEM from independent biological replicates (n=3). ns-not significant, * P<0.05, ** P<0.01, *** P<0.001, **** P < 0.0001; unpaired t test

Amplified expression of PDGFRA has been found in 9.9% of LGG, and was more frequently observed in diffuse astrocytoma (16.3%) as compared to oligodendroglioma (2.6%). (43) We developed a second mIDH1 mouse glioma model using the PDGFRA^D842V^ plasmid to constitutively activate the receptor tyrosine kinase (RTK)-RAS-PI3K pathway (Supplemental Figure 6A). We induced brain tumors in two experimental groups: wtIDH1 (RPA: PDGFRA^D842V^, shP53, and shATRX) and mIDH1 (RPAI: PDGFRA^D842V^, shP53, shATRX, and mIDH1^R132H^). The MS of mice in the RPAI group was 172 days post-injection (DPI), which was significantly greater compared to RPA group with an MS of 92 DPI; *P<0.0023*, Mantel-Cox test (Supplemental Figure 6B).

We generated the RPAI NS from a resected sleeping-beauty tumor 158 DPI, and integration of oncogenic plasmids was assessed by fluorescence microscopy (Supplemental Figure 6C-D). Additionally, we demonstrated that the RPAI NS was able to generate tumors in mice and confirmed expression of mIDH1 using a specific antibody for the R132H mutation and ATRX loss by IHC (Supplemental Figure 6E). WB analysis of RPAI NS treated with AGI-5198 revealed a decrease in global histone mark lysine trimethylation for H3K4me3, H3K27me3 and H3K36me3, but there was no change in global H3K27ac expression (Supplemental Figure 6F). After we confirmed the loss of ZMYND8 in RPAI ZMYND8 KO by WB analysis (Supplemental Figure 6G), we assessed their viability in response to IR. The RPAI ZMYND8 KO exhibited an IC_50_ of 4.0Gy versus the RPAI ZMYND8 WT with an IC_50_ of 8.1Gy (Supplemental Figure 6H). This finding further supports the importance of ZMYND8 in survival of mIDH1-expression glioma cells to IR independent of the oncogenic driver mutation.

To begin to examine the mechanisms by which ZMYND8 contributes to genomic stability, we inhibited two newly described ZMYND8-interacting partners BRD4 and HDAC. We sought to investigate if BRD4 and HDAC cooperate with ZMYND8 to mediate the response to irradiation. We evaluated the sensitivity of our two mouse mIDH1 NS, NPAI ZMYND8 WT versus NPAI ZMYND8 KO and RPAI ZMYND8 WT versus RPAI ZMYND8 KO to two bromodomain and extraterminal domain inhibitors (BETi): JQ1 and I-BET-762, which also target BRD2 and BRD3 in addition to BRD4 (Figure 4H). We also assessed the sensitivity of our mouse mIDH1 NS to the pan-histone deacetylase inhibitor Panobinostat, which targets Class I, II and IV HDACs (Figure 4I). We observed an enhanced vulnerability of NPAI ZMYND8 KO to JQ1 treatment (*P > 0.0001*) with an IC_50_ of 0.18µM versus the NPAI ZMYND8 WT where an IC_50_ was not reached (Supplemental Figure 7A). The RPAI ZMYND8 KO had a reduced viability to JQ1 (*P > 0.01*) with an IC_50_ of 0.51µM versus the RPAI ZMYND8 WT with an IC_50_ of 0.97µM (Supplemental Figure 7B). The NPAI ZMYND8 KO displayed reduced cellular viability to I-BET-762 (*P > 0.0001*) with an IC_50_ of 0.335µM versus the NPAI ZMYND8 WT where an IC_50_ was not reached (Supplemental Figure 7C). The RPAI ZMYND8 KO displayed reduced viability to I-BET-762 (*P > 0.0001*) with an IC_50_ of 0.001µM versus the RPAI ZMYND8 WT with an IC_50_ of 0.19µM (Supplemental Figure 7D). Both mouse NPAI NS displayed sensitivity to Panobinostat (*P > 0.0001*), with NPAI ZMYND8 WT having an IC_50_ of 0.0340µM versus NPAI ZMYND8 KO with an IC_50_ of 0.0075µM (Supplemental Figure 7E). Similarly, both RPAI NS showed sensitivity to Panobinostat (*P > 0.0001*), with RPAI ZMYND8 KO having an IC_50_ of 0.0082µM versus RPAI ZMYND8 WT with an IC_50_ of 0.41µM (Supplemental Figure 7F). We found that the NPAI ZMYND8 KO were more sensitive to BRD4 and HDAC inhibition compared to the NPAI ZMYND8 WT, which was also confirmed in the RPAI ZMYND8 KO versus RPAI ZMYND8 WT. The ZMYND8 KO mIDH1 GCCs exhibited higher levels of radiosensitization when IR was delivered in combination with either BRD4 or HDAC inhibition (Figure 4J). Collectively, this data suggests the inhibition of epigenetic modulators BRD4 or HDAC in combination with IR further reduced the cellular viability of ZMYND8 KO mIDH1 GCCs.

### ZMYND8 KO mIDH1 GCCs are defective in resolving IR induced DNA damage and undergo activation of cell cycle arrest

To further examine the relationship between ZMYND8 and DDR signaling, we analyzed the activation of HR proteins at the indicated time points following 20Gy IR in SF10602 ZMYND8 WT and SF10602 ZMYND8 KO cells by WB (Figure 5A). The HR pathway maintains genomic integrity by first sensing DNA damage, recruiting DNA repair mediators to the region and inducing cell cycle arrest (Figure 5B). Gamma histone H2AX (γH2AX) has been used to assess DNA damage following IR, but can also indicate genomic instability in the form of replication stress. (44, 45) We validated the loss of ZMYND8 expression in the SF10602 ZMYND8 KO (Figure 5C) and quantified the changes in ZMYND8 expression within the SF10602 ZMYND8 WT post-IR (Figure 5D). We observed sustained γH2AX signal in SF10602 ZMYND8 KO compared with SF10602 ZMYND8 WT following IR (Figure 5C). SF10602 ZMYND8 KO display higher basal γH2AX signal in the non-irradiated (NR) control when compared with SF10602 ZMYND8 WT (Figure 5E). The greatest difference in γH2AX expression relative to loading control (tubulin) was present at 24hrs post-IR (*P < 0.001*) (Figure 5E). This delayed resolution of γH2AX signal may reflect a defect in the efficiency of DSB repair or an accumulation of DNA damage in the SF10602 ZMYND8 KO following IR. ATM is a master regulator of HR repair and undergoes autophosphorylation at serine 1981 (pATM), which actives ATM by dissociating it from a dimer to a monomer form. (46) We observed enhanced activation of pATM in the SF10602 ZMYND8 KO compared to SF10602 ZMYND8 WT in response to IR (Figure 5F). The greatest difference in pATM activation relative to β-actin was present at 30mins post-IR (*P < 0.001*) (Figure 5G). DNA damage acquired during IR interrupts cell cycle progression through the activation of checkpoint kinases Chk1 and Chk2 to allow accurate DNA repair fidelity. As a safeguard, Chk1 triggers G2/M arrest in response to IR-induced DNA damage, its phosphorylation at serine 345 (pChk1) by ATR signifies impaired replication control. (47) We observed extended activation of pChk1 relative to Chkl in SF10602 ZMYND8 KO compared to SF10602 ZMYND8 WT in response to IR (Figure 5F). While the SF10602 ZMYND8 WT show a return of pChk1 to NR levels at 48hrs post-IR (*P < 0.0001*), the SF10602 ZMYND8 KO display prolonged activation of pChk1 (Figure 5H).

**Figure 5:**
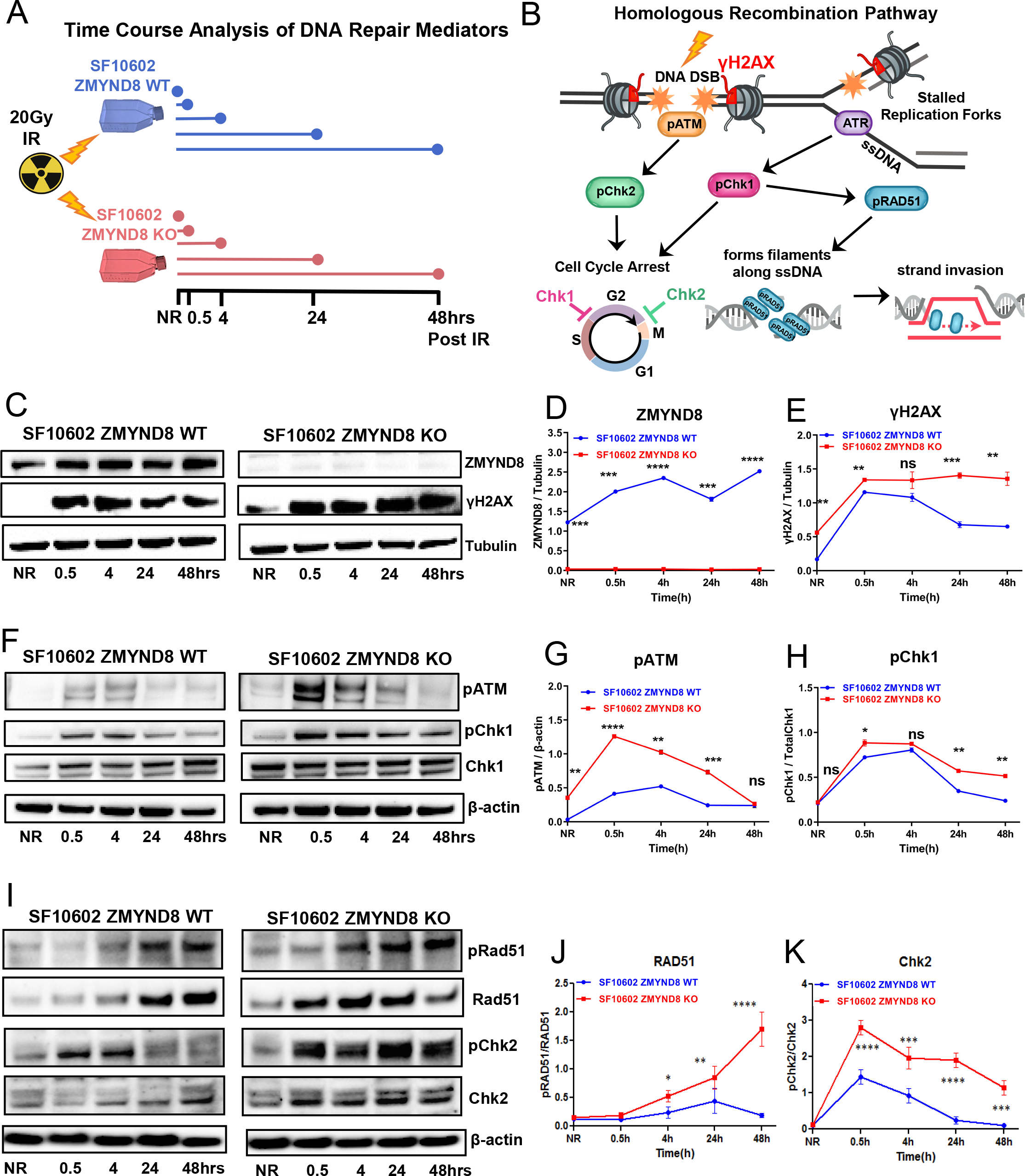
ZMYND8 KO mIDH1 GCCs are defective in resolving IR induced DNA damage and undergo prolonged activation of cell cycle arrest. **(A**). Diagram of the time course analysis of DNA repair proteins expressed in SF10602 ZMYND8 WT (blue) vs. SF10602 ZMYND8 KO (red) exposed to a single dose of 20Gy IR and protein was collected from non-irradiated (NR) cellls and at 0.5, 4, 24, and 48 hours post IR exposure. (**B**) Model of irradiation (lighting blot) induced activation of the homologous recombination (HR) pathway including downstream HR mediators. (**C**) Representative western blot for ZMYND8 and γH2AX expression in SF10602 ZMYND8 WT vs.SF10602 ZMYND8 KO at the indicated conditions described in IR time course diagram. (**D**) Line graph represents the quantification of ZMYND8 expression relative to tubulin in SF10602 ZMYND8 WT (blue) vs. SF10602 KO (red) from 0 (NR) to 48hrs post IR. (**E**) Line graph represents the quantification of yH2AX relative to tubulin. (**F**) Representative western blot for phosphorylated ATM (pATM) and Chk1 (pChk1) expression and their respective nonphosphorylated (ATM and Chk1) proteins relative to β-actin. Line graph represents the quantification of (**G**) pATM expression relative to β-actin and (**H**) pChk1 expression relative to Chk1. (**I**) Representative western blot for phosphorylated Rad51 (pRad51) and Chk2 (pChk2) expression and their respective nonphosphorylated (Rad51 and Chk2) proteins relative to β-actin. Line graph represents the quantification of (**J**) pRad51 expression relative to Rad51 and (**H**) pChk2 expression relative to Chk2. Error bars represent SEM from independent biological replicates. (n=3) * P<0.05, ** P<0.01, *** P<0.001, **** P < 0.0001; unpaired t test.

Next, we evaluated phosphorylation of Rad51 at threonine 309 (pRad51), which is mediated by activated Chk1. Rad51 promotes genomic stability by binding to ssDNA to stabilize replication forks in order to protect under-replicated DNA regions during mitosis. (48–50) Additionally, Rad51 is recruited to DSB regions where it cooperates with HR proteins to promote strand invasion. (48) We observed elevated relative expression of pRad/Rad51 in SF10602 ZMYND8 KO compared to SF10602 ZMYND8 WT beginning at 4hrs post-IR (*P < 0.05)* which was most significantly elevated after 48hrs post IR (*P < 0.0001)* (Figure 5I-J). Chk2 is phosphorylated on threonine 68 (pChk2) by ATM in order to induce G2/M arrest. We observed significant increase in the relative pChk2/Chk2 level in the SF10602 ZMYND8 KO compared to the SF10602 ZMYND8 WT beginning at 30mins post-IR (*P < 0.0001)* (Figure 5K). While the SF10602 ZMYND8 WT show a return of pChk2 to NR levels at 24hrs post-IR (*P < 0.0001*), the SF10602 ZMYND8 KO display prolonged activation of pChk2 (Figure 5K). These findings suggest that ZMYND8 plays an important role in DDR, which is mediated by the mobilization of DDR proteins to repair IR-induced DNA damage and regulation of cell cycle progression.

### ZMYND8 KO mIDH1 GCCs are more susceptible to PARP inhibition

Considering the elevated expression of DDR proteins following IR in ZMYND8 KO GCCs compared to ZMYND8 WT GCCs, we wondered if this resulted from an impaired resolution of DNA damage due to intrinsic genomic stress occurring within the ZMYND8 KO GCCs. We speculated that loss of ZMYND8 would increase the susceptibility of mIDH1 GCCs to chemicals that promote genotoxic stress. To evaluate this, we targeted PARP, an initial sensor of DNA lesions specifically at single-stranded DNA breaks (SSB) (Figure 6A). PARP1 and PARP2 catalyze a reaction that utilizes NAD+ to attach negatively charged poly(ADP-ribose) polymers to various target proteins, including itself, to signal the recruitment of DNA repair proteins to the region. We choose to utilize pamiparib, a selective PARP1/2 inhibitor that traps PARP to DNA forming PARP-DNA complexes that allow for the persistence of unrepaired SSB (Figure 6A). (51) The PARP-DNA adducts create a barrier at replication forks, resulting in fork collapse and the formation of DSBs (Figure 6A). (51) We observed a significant reduction (*P < 0.001*) in cellular viability of the SF10602 ZMYND8 KO to increasing doses of pamiparib with an IC_50_ of 12.6µM versus the SF10602 ZMYND8 WT with an IC_50_ of 88.5 µM (Figure 6B). When we combined Pamiparib with 20Gy IR, we observed enhanced therapeutic response in the SF10602 ZMYND8 KO compared to the SF10602 ZMYND8 WT (Figure 6C). The sensitivity of SF10602 to Pamiparib +IR was further enhanced by the loss of ZMYND8, where this combination was also supported in the MGG119 (Supplemental Figure S8A-B). Next, we evaluated the response of our mouse mIDH1 GCCs to PARP inhibition; we observed a significant reduction in cellular viability of NPAI ZMYND8 KO to Pamiparib alone with an IC_50_ of 4.9µM versus the NPAI ZMYND8 WT with an IC_50_ of 50.5µM (Figure 6D). Comparably, the RPAI ZMYND8 KO displayed a significant reduction in cellular viability to Pamiparib alone (*P > 0.0001*) with an IC_50_ of 2.0µM versus the NPAI ZMYND8 WT with an IC_50_ of 44.7µM (Supplemental Figure S8C). In the combination of Pamiparib and 6Gy IR, we similarly observed an enhanced therapeutic response in the NPAI ZMYND8 KO compared to the NPAI ZMYND8 WT (Figure 6E). Additionally, the combination of Pamiparib and IR greatly reduced the cellular viability of RPAI ZMYND8 KO compared to RPAI ZMYND8 WT (Supplemental Figure S8D). To translate these findings to *in vivo*, we implanted 50,000 NPAI ZMYND8 WT GCCs in immunocompetent C57BL6 mice and confirmed tumor burden based on IVIS bioluminescence signal at 10 days post implantation (dpi). Tumor bearing mice were administered saline or pamiparib (1mg/mL) and with/out IR, as indicated in Figure 6F. We observed a 1.7-fold (*P < 0.05*) increase in the MS of mice in the Pamiparib + IR group (MS: 55dpi) when compared to IR alone (MS: 38 dpi) or Pamiparib alone (MS:34 dpi) (Figure 6G). When compared to the control mice that received saline (MS:29 dpi), mice in the Pampiparib + IR group displayed a 1.9-fold (*P<0.01*) increase in MS (Figure 6G). We did not observe a significant difference in viability in response to AGI-5198 treatment in NPAI ZMYND8 WT versus ZMYND8 KO, but when combined with IR we observed a modest reduction in viability *P<0.05* (Figure 6H). The enhanced sensitivity of mIDH1 GCCs to IR after the genetic knockout of ZMYND8, or treatment with HDAC or PARP inhibitors, propose a novel vulnerability of mIDH1 GCCs to respond to IR-induced DNA damage. We speculate that the loss of transcriptional repression mediated by ZMYND8’s recruitment of NuRD or ZMYND8 recognition of DNA damaged regions marked by PARP leads to defective HR repair in mIDH1 GCCs (Figure 6I).

**Figure 6:**
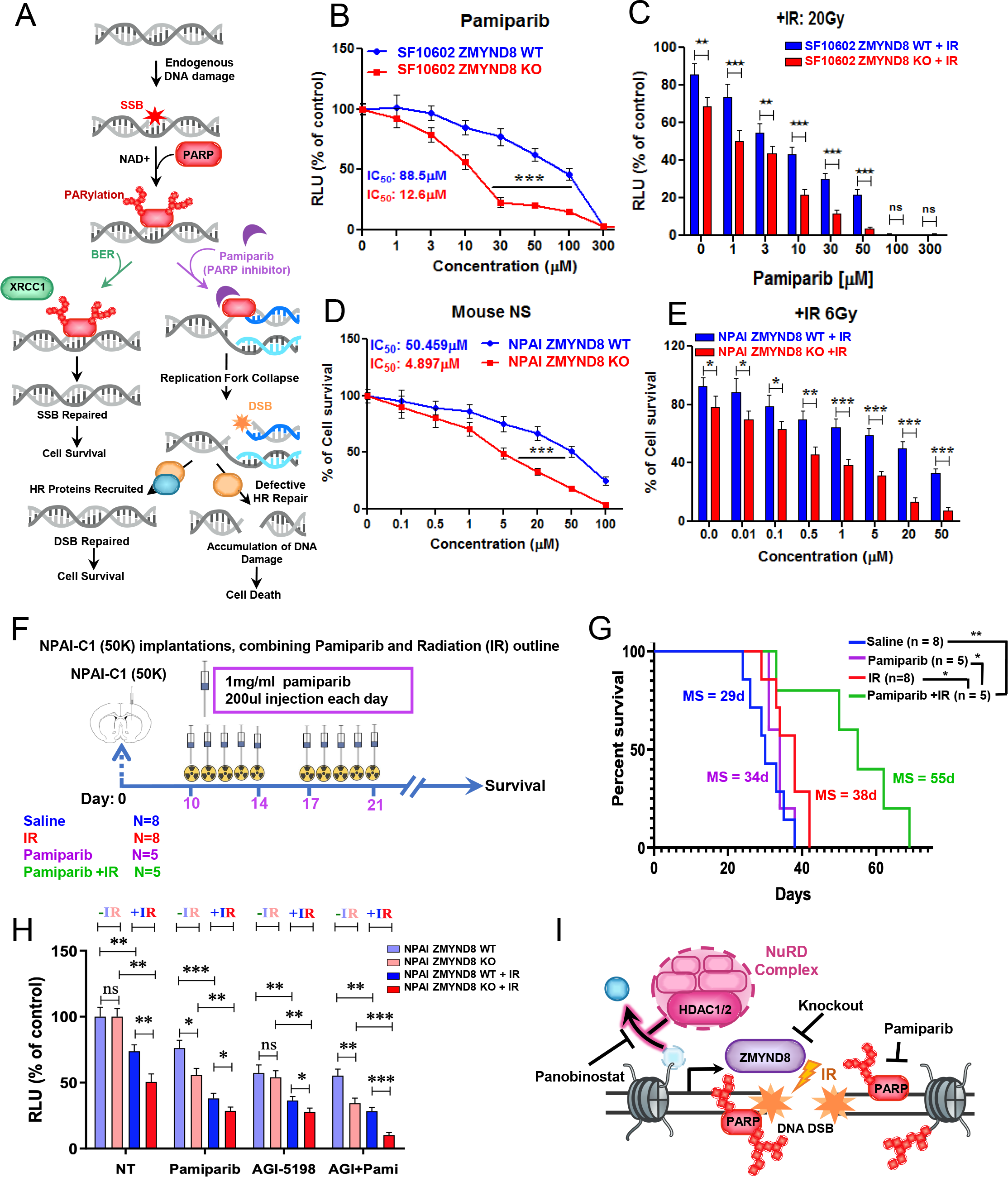
ZMYND8 KO mIDH1 GCCs are more susceptible to PARP inhibition. (A) Proposed mechanism of PARP’s repair of single stranded breaks (SSB), which occur as a results of endogenous DNA damage in proliferating cells. PARP1/2 catalyze a reaction that utilizes nicotinamide adenine dinucleotide (NAD+) to add poly(ADP-ribose) polymers to itself and signals the recruitment of proteins involved in base excision repair (BER) like XRCC1 to resolve the SSB. Pamiparib (PARP inhibitor) block PARylation and trap PARP onto DNA, resulting in the formation of double stranded breaks (DSB) at collapsed replication forks. Cells that can successfully recruit HR proteins can repair the DSB. (**B**) Pamiparib dose response curve to evaluate the impact of PARP inhibition on SF10602 ZMYND8 WT (blue) vs. SF10602 ZMYND8 KO (red). Two-tailed t-test (**C**) Cell viability assay shows the effect of Pamiparib + Irradiation (IR 20Gy) on cell proliferation in SF10602 ZMYND8 WT vs. SF10602 ZMYND8 KO measured in RLU relative to control non-treated. Two-tailed t-test (**D**) Pamiparib dose response curve from mouse NPAI mIDH1 GCCs ZMYND8 WT (blue) vs. NPAI ZMYND8 KO (red). Two-tailed t-test (**E**) Cell viability assay shows the effect of Pamiparib + IR 6Gy on cell proliferation in NPAI ZMYND8 WT (blue) vs. NPAI ZMYND8 KO (red) measured in RLU relative to control IR alone (0μM Pamiparib). Two-tailed t-test (**F**) Preclinical design for testing the impact of PARP inhibitor (Pamiparib) on the response to IR in an orthotopic glioma model. Ten days after implantation of 50,000 NPAI NS, animals were randomly divided into 4 groups: (i) saline, (ii) IR received 2Gy/day for 5 days each cycle (20Gy total) starting 10 days post implantation (dpi) (iii) pamiparib delivered intraperitoneal (1mg/mL injection) 5 days on and 2 days off for 2 weeks, (iv) pamiparib + IR. Mice were monitored for tumor burden and euthanized at symptomatic stages to track survival to treatment. (**G**) Kaplan-Meier survival curve of NPAI ZMYND8 WT tumor bearing mice treated with or without 6Gy (n=8) in the presence or absence of Pamiparib (n=5). \**P* < 0.05, \*\**P* < 0.01; log-rank Mantel-Cox test. (**H**) Impact of mIDH1 inhibition (AGI-5198) and Pamiparib alone or in combination on the cellular viability of NPAI ZMYND8 WT in the presence (+IR) or absence (-IR) of irradiation. Errors bars represent SEM from independent biological replicates (n=3). ns-not significant, * P<0.05, ** P<0.01, *** P<0.001, **** P < 0.0001; Multiple t-test was used (**I**) Working model in which targeting ZMYND8 by genetic knockout or treatment with HDAC or PARP inhibition exposes a epigenetic vulnerability of mIDH1 GCCs to IR induced DNA damage.

To address whether ZMYND8 expression varied across LGG patients based on IDH1 mutation status, we analyzed publically available RNA-seq data from the The Cancer Genome Atlas (TCGA) LGG dataset (Supplemental Figure S9A). We observed a significant increase (*P<0.05*) in log2-normalized ZMYND8 expression in mIDH1 LGG versus wtIDH1 LGG. When we stratified the overall survival (OS) of LGG patient into low- or high-ZMYND8 expression groups, we found that high-ZMYND8 mIDH1 LGG were associated with poor clinical outcome with a median OS of 6.3yrs as compared with low-ZMYND8 mIDH1 LGG with a median OS of 7.9yrs (Supplemental Figure S9B). There was no significant difference found within wtIDH1 LGGs, where high-ZMYND8 wtIDH1 LGGs had a median OS of 1.7yrs compared with 2.0yrs in low-ZMYND8 wtIDH1 LGGs. This significant discrepancy in MS found amongst mIDH1 LGGs (*P<0.0001*), indicates that a subset of mIDH1 glioma overexpress ZMYND8 and of which present with poor patient outcome.

## Discussion

IDH-mutant gliomas represent a distinct molecular subtype of brain cancer defined by the presence of DNA hypermethylation commonly referred to as CpG island methylator phenotype (CIMP) and dysregulation in the removal of histone lysine trimethylation. (7, 8) This epigenetic reprogramming mechanism is dependent upon the production of 2HG by mIDH1, which notably our group and others have demonstrated can be reversed through treatment with mIDH1 specific inhibitors. (9,10,28,52) A shared feature across IDH1/2 mutant cancers showed a high propensity of DNA hypermethylated regions at gene bodies and enhancers, while surprisingly the majority of gene promoters remained hypomethylated. (52) Although this finding might imply a universal effect of the IDH-mutation, another study revealed distinct methylome patterns when comparing IDH- mutant tumors to IDH-wildtype tissue matched counterparts. The authors found that 19% of CpG sites were exclusively hypermethylated in IDH-mutant glioma compared to 3% in IDH-mutant AML, 2% in melanoma, and 4% in cholangiocarcinoma. (53) CpG methylation can impact nearby genes altering their expression, thus these differentially methylated regions across IDH-mutant cancers may be dependent upon the cellular origin of the cancer. IDH-mutant gliomas displayed the greatest number of biological pathway alterations compared with the other tumor types, specifically the downregulation of tissue development and an upregulation of a subset of DNA repair pathways. (53)

To date, our understanding of the reversibility of histone hypermethylation following loss of mIDH1 activity has been limited to immortalized human astrocytes (IHA) that expressed mIDH1 under a doxycycline (dox) inducible system. (7) Notably, IHA-expressing mIDH1 displayed upregulation of stem-like genes: *CD24* and *PDGFRA*, which was accompanied by a transient gain of H3K4me3 at these gene promoter regions suggesting that mIDH1 poises these genes for activation. Following dox-withdrawal, methylation at most loci eventually returned to baseline levels either transiently (13.8%) or gradually (62.5%) across cell passages, but 23.7% of regions persisted after loss of mIDH1. These findings suggest that the epigenetic reprogramming mediated by mIDH1 is still not well understood, especially in regard to the contribution of mIDH1 regulation on genes involved in therapeutic resistance to radiation treatment.

Given that mIDH1-R132H is endogenously expressed in SF10602 and was derived from a patient tumor subjected to radiotherapy, we sought to determine mIDH1-dependent mechanisms that contributed to radioresistance. (27) We found that ZMYND8 was significantly downregulated following mIDH1 inhibition in the SF10602 treated with AGI-5198. Recent studies have reported that ZMYND8’s recruitment to regions of laser-microirradation induced DNA damage is important to facilitate HR DNA repair. (12,14,15) In doing so, ZMYND8 functions as a beacon to direct the NuRD complex to damaged DNA regions and allow for repression of transcription. (12, 15) The acetylated histone residues H3K14ac and H4K16ac, which are recognized by ZMYND8, have been shown to be crucial for DNA checkpoint activation and local HR repair of DSB. (54, 55) Recent studies have demonstrated preferential vulnerability of mIDH1-expressing glioma models to inhibitors targeting HDAC, which is a component of the NuRD complex. (56, 57) Interestingly, mIDH1 glioma cells treated with panobinostat displayed increased H3K14ac, which is recognized by ZMYND8. (57)

Our current findings provide evidence that ZMYND8 is epigenetically regulated in human mIDH1 GCCs based on the loss of active histone mark H3K4me3 at the ZMYND8 promoter locus following mIDH1 inhibition. Additionally, our comparison of mouse mIDH1 NS vs. wtIDH1 NS demonstrated that mouse mIDH1 NS had an enhanced enrichment of H3K4me3 at the ZMYND8 promoter locus by ChIP-seq and increased nascent transcription by Bru-seq. To our knowledge, a direct connection between ZMYND8 regulation at the chromatin level within mIDH1 glioma has not previously been described. However, we found that high ZMYND8 expressing mIDH1 LGG patient tumors exhibited a shorter median OS as compared with low ZMYND8 expressing mIDH1 LGG. In the context of brain cancer, ZMYND8 has been shown to preferentially bind histone 3.3 point mutant G34R (H3.3G34R) present in pediatric high grade glioma to suppress genes involved in MHC II presentation. (21)

We observed a significant reduction in ZMYND8 protein expression in three patient derived mIDH1 GCC after treatment with DS-1001b. Additionally, inhibition of mIDH1 in SF10602 increased expression of genes associated with replication stress and genomic instability. Timeless has been shown to accumulate at regions of DNA damage tracks induced by laser-microirradiation and its retention at damaged chromatin is dependent on the presence of PARP but not its activity. (58, 59) Cancer cells undergo persistent DNA replication stress as a result of aberrant cell cycle progression. Timeless has been shown to stabilize replication forks and contribute to sister chromatid cohesion for maintenance of genomic integrity. (60, 61) In cervical cancer models, the overexpression of Timeless has been proposed to function in response to replication stress driven by oncogene activation. (62) TREX1 is known to function as a 3’-5’ DNA exonuclease to degrade ssDNA or mispaired DNA duplexes that arise from the repair of DNA lesions and aberrant replication. (35, 49) The elevated expression of Timeless and TREX1 following mIDH1 inhibition in SF10602 could be an indication of enhanced genomic stress.

By selectively suppressing ZMYND8 expression in human mIDH1 GCCs by the means of shRNA knockdown or genetic knockout using lenti-CRISPRCas9, we observed a significant decrease in cellular viability in response to IR. Additionally, inhibition of epigenetic regulators BRD4 and HDAC further reduced the viability of ZMYND8 KO mIDH1 GCCs. Recent work in AML showed that ZMYND8 recruitment to enhancers was mediated by BRD4. (22) The oncogenic programs regulated by BRD4 to support cancer cell survival were downregulated in AMLs when ZMYND8 was depleted. (22) Inhibition of BRD4 using JQ1 was shown to abolish ZMYND8 occupancy at enhancers. (22) There is compelling evidence that BRD4 functions at replication forks, where it associates with several complexes involved in chromatin remodeling (MMS22L, TONSL) and replication (POLA2, POLD3), which we also observed to be upregulated in our RNA-seq analysis. (63) Additionally, inhibition of BRD4 either by siRNA knockdown or JQ1 treatment has been shown to induce DNA damage marked by increased γH2AX foci formation. (64) Loss of BRD4 induces replicative stress via the deregulation of transcription and in response RAD51 is recruited to damaged DNA regions to suppress replication. (65) Treatment of mIDH1 tumor cells with the HDAC inhibitor vorinostat lead to a suppression of Rad51 and Brca1 expression, along with an increase in cleaved PARP. (56)

In support of the role of ZMYND8 in genomic integrity, loss of ZMYND8 in triple-negative breast cancer cells lead to an increase in micronuclei formation along with chromosome aberrations including dicentric chromosomes and DNA breaks evaluated by metaphase spreads. (20) A potential mechanism contributing to this induction of genomic instability, could be the result of overactivation of enhancers following the loss of ZMYND8. (54) A recent study mapping transcription-mediated DSB in breast cancer revealed that RAD51 localized to super-enhancers to stabilize DNA damage occurring due to hypertranscription. (66)

In our time-course assessment of HR repair protein activation following IR, SF10602 ZMYND8 KO displayed prolonged activation of pChk2, while the SF10602 ZMYND8 WT showed a return to baseline 24hrs post-IR. We hypothesize that loss of ZMYND8 impairs the resolution of DSB induced by IR, which leads to extended cell cycle arrest mediated by pChk2.

PARP functions as an early sensor of DNA damage and has been shown to rapidly deposit poly-ADP ribose chains (PARylation) at newly generated DNA DSB within milliseconds. (67)The recruitment of ZMYND8 to laser-induced DSB was shown to be dependent upon the presence of PARylation. (15)Surprisingly, we observed that ZMYND8 WT mIDH1 glioma cells were sensitive to Pamiparib treatment, but ZMYND8 KO mIDH1 glioma cells displayed a further decrease in cellular viability. Recent studies have observed enhanced sensitivity of IDH-mutant glioma to PARP inhibition using preclinical mouse models and human glioma stem cell lines. (68, 69) Pamiparib (BGB-290) in combination with TMZ is currently being evaluated clinically for the treatment of IDH1/2 mutant grade I-IV Gliomas (NCT03749187) and recurrent gliomas with IDH1/2 mutations (NCT03914742). Moreover, our data shows that ZMYND8 functions alongside PARP to signal the repair of IR induced DNA damage.

In summary, by examining the alterations to the epigenome and biological pathways following mIDH1 inhibition, we provide novel evidence that ZMYND8 is epigenetically regulated by mIDH1 reprogramming in patient derived mIDH1 GCCs and in our genetically engineered mouse mIDH1 glioma model. In addition, our data shows that ZMYND8 supports the survival of human mIDH1 GCCs and mouse mIDH1 NS in response to radiation through its role in contributing to DNA repair and genomic stability. Genetic ablation of ZMYND8 in combination with epigenetic inhibitors against PARP, HDAC, and BRD4 further reduces the cellular viability of mIDH1 GCCs to IR induced DNA damage. We anticipate these findings will provide support for the development of ZMYND8 specific inhibitors, which represents a viable target for the treatment of mIDH1 gliomas.

## Acknowledgments

We thank D. Cahill (Harvard Medical School) for providing the mIDH1 human glioma cells MGG119 and Dr. J. Costello (UCSF) for providing the mIDH1 human glioma cells SF10602; Dr. M. Luo and Dr. W. Lou for providing the lentiviral ZMYND8 CRISPRv2 plasmid; and Dr. J. Ohlfest (University of Minnesota, deceased) for providing the SB model plasmids.

