## Supplemental Materials for "Zinc Finger MYND-Type Containing 8 (ZMYND8) is epigenetically regulated in mutant Isocitrate Dehydrogenase 1 (IDH1) glioma to promote radioresistance"

This PDF file includes:

Supplementary Methods and References

Supplementary Figures S1-S9

Supplementary Table S1

#### Supplementary methods

##### Bru-seq and BruChase-seq

These techniques were performed as described previously (*Paulsen et al. 2014*). Briefly, nascent RNA labelling was performed for 30 min at 37 °C with 2 mM 5-bromouridine (Sigma-Aldrich, 850187) in conditioned medium. For the BruChase-seq analysis, the bromouridine-containing medium was removed after labelling for 30 min, the plates were rinsed twice in PBS, and then conditioned medium containing 20 mM uridine (Roche, 11934554001) was added. The cells were then incubated for 6 hours at 37 °C. At the completion of labelling with or without chase, the cells were lysed in TRIzol (Thermo Fisher Scientific, 15596018), and the Bru-labeled RNA was captured with anti-BrdU antibodies (BD Biosciences, 555627) conjugated to magnetic beads (Dynabeads goat anti-mouse IgG, Thermo Fisher Scientific, 11033). The isolated RNA was converted into cDNA libraries and prepared for sequencing using the Illumina TruSeq RNA Library Preparation Kit v2 (Illumina, RS-122-2001) followed by deep sequencing to around 50 million single-end 50 nucleotide reads.

##### Bru-seq data analysis

Bioinformatics and data analysis pipeline was implemented using the q pipeline manager (http://sourceforge.net/projects/qppln-mngr/). The bioinformatics programs used, read mapping, genome annotation, and expression scoring were previously described (*Paulsen et al. 2014*), section 2.4. (1) RPKM (reads per kilobase per million mapped reads) values were calculated for individual genes that were at least 300 bp long. For genes with lengths of 30 kb and less, RPKM values were calculated using read counts from the entire gene. For genes longer than 30 kb, an RPKM value was calculated using read counts from the first 30 kb downstream of the TSS. The R package DESeq35 was used to test differential expression of genes whose mean RPKM between samples was greater than 0.5. Significant changes in transcription initiation were defined as follows: adjusted p-values < 0.05; fold change < 1.5.

**Quantitative Pathology & Bioimage Analysis Software Analysis**

QuPath v0.3.2(actively developed at the University of Edinburgh) is open-source software for bioimage analysis and digital pathology. (2) Positive cell detection command allows to classify cells as either positive or negative, using DAB optical density mean and a single threshold. This command allows to detect 'objects'(cells) in the selected field, and the percentage of representation of each cluster (negative and positive, in this case). We utilized 13-15 frames at 20X magnification consisting of 3-5 distinct neutrosphere sections for each frame. These same parameters were employed for the detection of all the fields for every sample condition. Percentage of DAB-stained positive cells over total number of cells per fields was quantified. (Threshold: SF10602: 0.2; LC1035: 0.3).

**pT2-shZMYND8-GFP plasmid construction**

These plasmids were generated by excising the micro-30a loop from the pT2-shATRX-GFP (Addgene #124259) via overnight restriction enzyme digestion with XhoI and EcoRI at 37ºC. The backbone pT2-GFP DNA was purified using the QIAquick Gel Extraction Kit (Cat No: 28706). Oligonucleotide sequences consisting of short hairpin (sh) that target exon 12 of the human ZMYND8 gene or exon 24 in the mouse ZYMND8 gene were ligated using T4 DNA ligase protocol (NEB M0202). The ligated pT2-shZMYND8-GFP plasmid was transformed into competent *E Coli* (NEB C2984H) and single colonies were expanded for DNA isolation using the QIAprep spin miniprep kit (Cat No: 27104). Cloned sites were confirmed by DNA sanger sequencing.

**Generation of ZMYND8 knockdown tumor cell lines**

Stable transfection of pT2-shZMYND8-GFP was performed using electroporation combined with sleeping-beauty transposon system integration *in vitro*. Human mIDH1 primary glioma cell cultures, SF10602 and MGG119 were dissociated to single cells and $1x10^6$ cells were collected. Glioma cell pellets were resuspended in Lonza Nucleofector (V4XP-3024) P3 buffer solution and nucleofected on the 4D-Nucleofector system (Cat No: AAF-1002B) along with pT2-shZMYND8-GFP (1.5 µg) and Sleeping beauty transpose-luciferase (Cat No.20207 Addgene, 0.5 µg) plasmids. Cells were transferred to laminin-coated plates and detection of green fluorescent cells was observed the next day by fluorescence microscopy. Isolation of GFP-positive glioma cells was performed 3-4 weeks after nucleofection in order to allow for higher enrichment. The expanded mIDH1 GCCs expressing shZMYND8 were sorted again to isolate the top 25% of GFP-expressing glioma cells. Knockdown of ZMYND8 expression was confirmed by WB.

**$PDGFR^{D842V}$-driven mouse mIDH1 glioma neurospheres**

We have generated an additional mIDH1 glioma model (RPAI) driven by platelet derived growth factor receptor alpha mutation ($PDGFRA^{D842V}$) to promote constituent activation of the MAPK pathway, which have been shown to induce tumors in diffuse intrinsic pontine glioma (DIPG) models. (3) These mouse glioma cells endogenous express IDH1-R132H, along with ATRX and TP53 short hairpin knockdown to simulate the molecular genetic lesions that define the molecular features of astrocytoma. Tumor NS were derived from endogenous mIDH1 tumors that were adapted to *in vitro* culture and could be reimplanted into NSG mice to demonstrate the ability to form tumors.

**Histone Extraction**

To assess changes in histone modifications in RPAI NS treated with either vehicle (DMSO) or mIDH1 inhibitor (AGI-5198), we collected histone extracts using the Histone Purification Mini

Kit (Active Motif, 40026). We performed WB analysis using 10µg histone protein and assessed global histone mark modifications for H3K4me3 (Diagenode, C15410003-50), H3K27me3 (Diagenode, C15410195), H3K27ac (Diagenode, C15410196) and H3K36me3 (Diagenode, C15410192) relative to total histone 3 (Cell Signaling, 9715S).

##### TCGA Analysis

Low Grade Glioma patient clinical annotation and RNA-seq data from the NCI Genomic Data Commons (GDC) repository were downloaded using the TCGAbiolinks package. (4) mRNA expression levels were normalized and ZMYND8 expression was compared between wtIDH1 and mIDH1 samples using a Log-rank test. The relationship between patients' overall survival and the levels of ZMYND8 were identified by the R packages "survminer" and "survival".

A

#### GSEA Enrichment Map of Differentially Expressed Gene Ontologies in mIDH1 inhibitor (AGI-5198) treated SF10602 compared to untreated SF10602

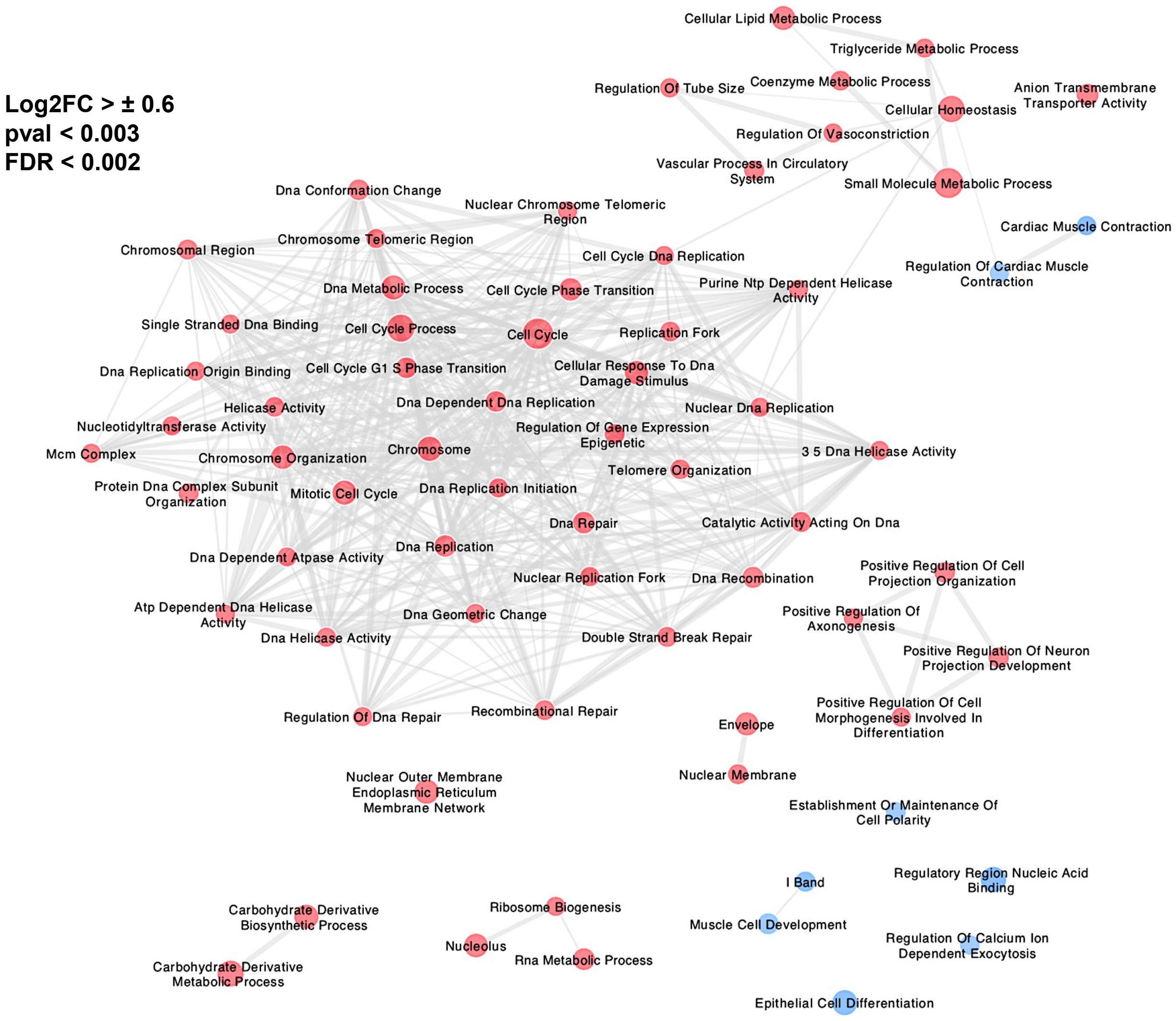

**Figure S1: Gene Set Enrichment Analysis (GSEA) Network Map of DE genes in mIDH1 inhibitor (AGI-5198) treated SF10602 mIDH1 GCC compared to untreated SF10602.** (**A**) GSEA pathway analysis of DE genes in AGI-5198 treated mIDH1 GCC SF10602 vs untreated. Circles represent distinct GOs and their sizes reflects the number of enriched genes within a GO. The cut-off used for defining DEG was log 2 fold-change(Log2C)>±0.6, p-value(pval)<0.003, false discovery rate(FDR)<0.002. Related GOs are linked by edges in gray for shared genes that have function within multiple pathways. The node color refers to the statistically significant DE genes in AGI-5198 treated vs untreated SF10602 where upregulated GOs are shown in red and downregulated GOs are shown in blue. Image was generated using Enrichment Map Plug-in on Cytoscape 3.9.1.

**Supplementary Fig S2**

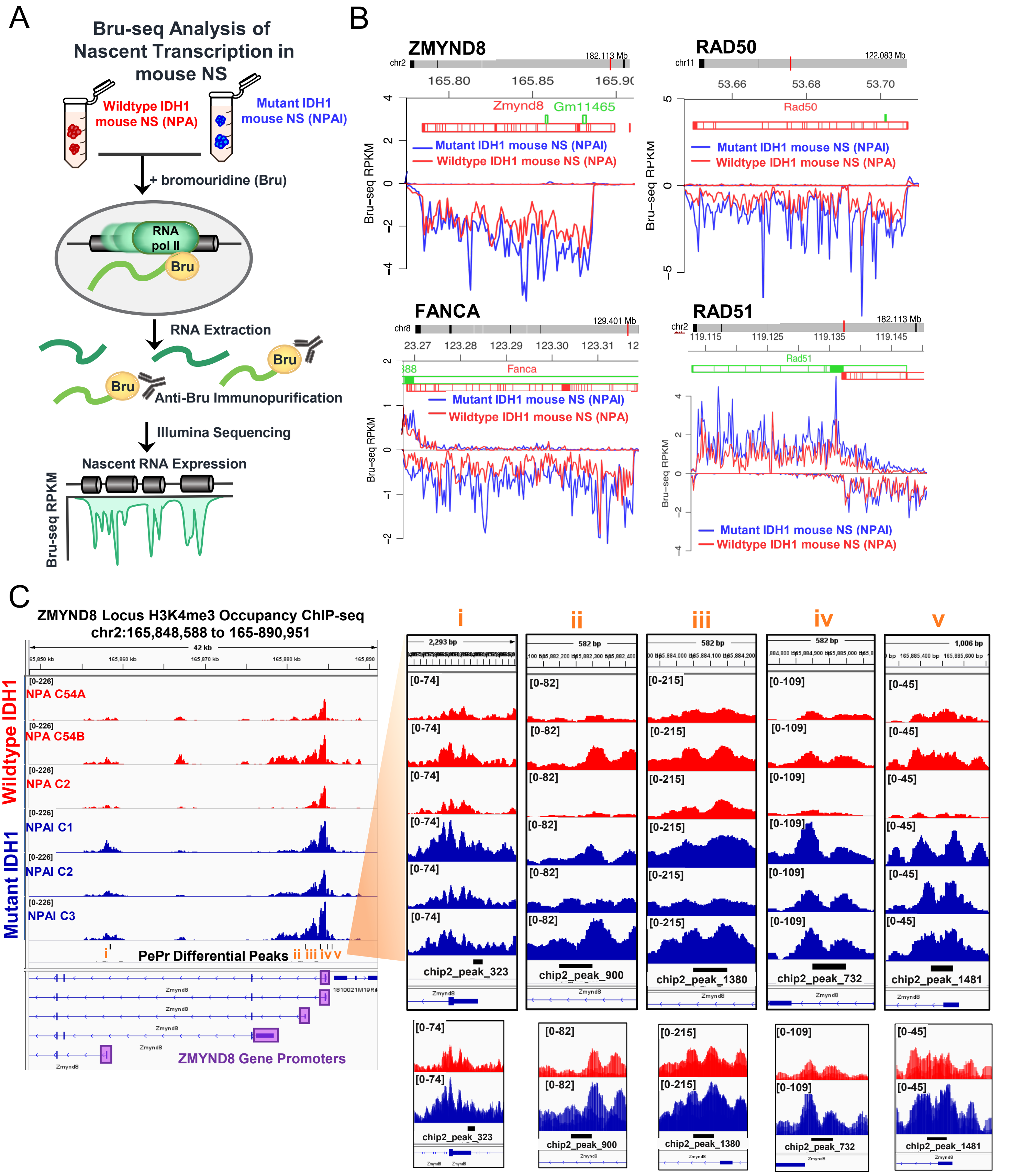

**Figure S2: Transcriptional and epigenetic regulation of the ZMYND8 locus in mutant IDH1 (mIDH1) vs. wildtype IDH1 (wtIDH1) mouse glioma model**

(**A**) Diagram of Bru-seq assay to quantify the rate of nascent transcription based on bromouridine labelled RNAs comparing wtIDH1 (red) vs. mIDH1 mouse NS (blue). **(B)** Bru-seq traces show differential transcriptional rates (<1.2fold ; P <0.05) of DNA repair genes ZMYND8, RAD50, FANCA and RAD51 in mIDH1 (blue, n=12) compared to wtIDH1 NS (red, n=3). **(C)** Integrative Genome Viewer (IGV) image displays ChIP-seq tracks of H3K4me3 enrichment at the ZMYND8 promoter region (purple) in wtIDH1 NS (red, n=3) vs. mIDH1 NS (blue, n=3). Differential enriched peaks (i-v, black bars) in mIDH1 NS vs wtIDH1 are displayed as separated tracks, while overlapped tracks are shown below. (*n* = 3 biological replicates per group).

### Supplementary Fig S3

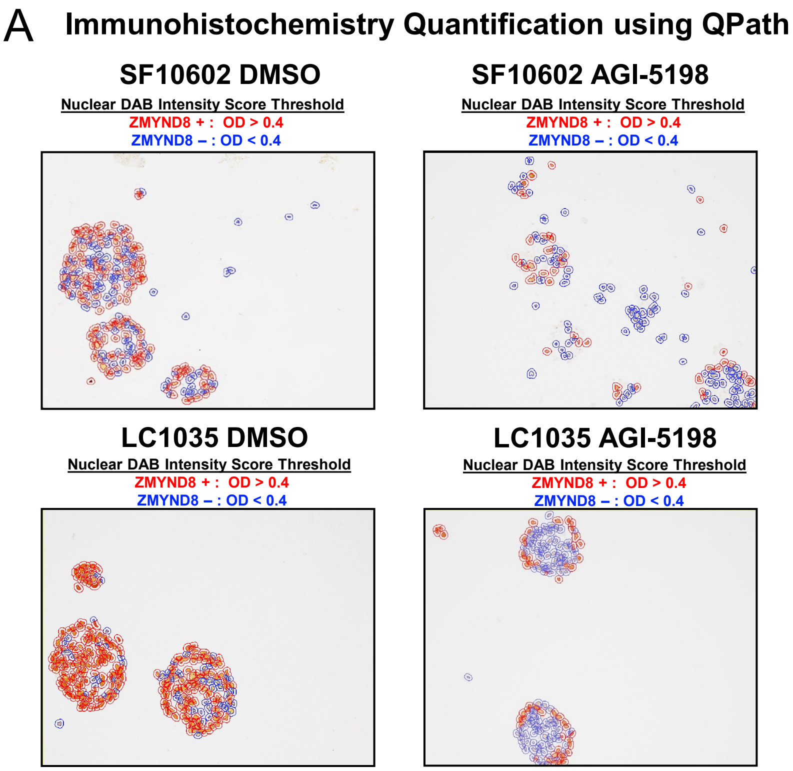


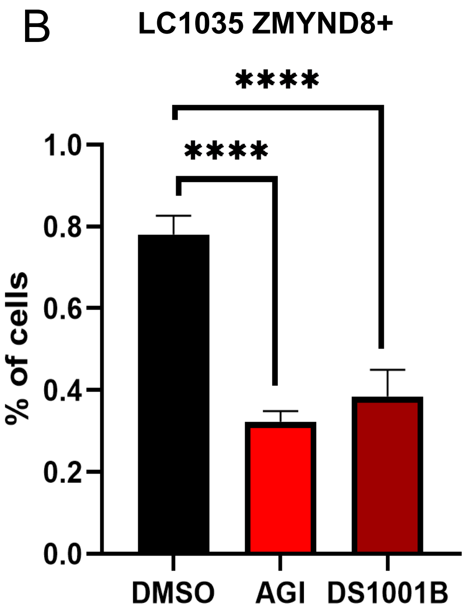

**Figure S3: Quantification of ZMYND8 positive mIDH1 GCCs in response to mIDH1 inhibitor treatment.**
(**A**) Representative QuPAth quantification images of nuclear ZMYND8 immunohistochemistry (IHC) staining in paraffin-embeded and sectioned human mIDH1 GCCs (SF10602,LC1035) treated with either vehicle (DMSO) or mIDH1 inhibitor (AGI-5198) for 1 week. Detection of ZMYND8 positive cells was based on a single cell DAB optical density (OD) threshold greater than 0.4 (red), while ZMYND8 negative cells had OD<0.4. **(B)** Histogram of the percentage of ZMYND8 positive cells counted in LC1035 treated with either DMSO, AGI-5198, or DS1001B for 1 week. Errors bars represent SEM from independent biological replicate frames (n=12). **** P < 0.0001; unpaired t test

### Supplementary Fig S4

A

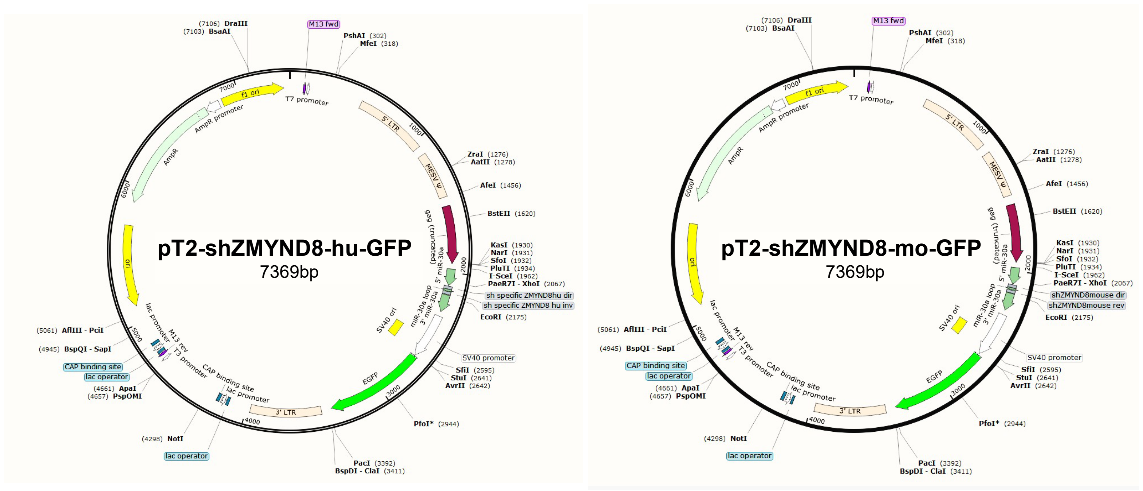


B

| shRNA | Sequences |
|---|---|
| Human sh2-ZMYND8 F | TCGAGAAGGTATATTGCTGTTGACAGTGAGCGCCGGATTTCCTTGTCGGATATTAGTGAAGCCACA GATGTAATATCCGACAAGGAAATCCGGTGCCTACTGCCTCGG |
| Human sh2-ZMYND8 R | AATTCCGAGGCAGTAGGCACCGGATTTCCTTGTCGGATATTACATCTGTGGCTTCACTAATATCCGA CAAGGAAATCCGGCGCTCACTGTCAACAGCAATATACCTTC |
| Mouse sh4-ZMYND8 F | TCGAGAAGGTATATTGCTGTTGACAGTGAGCGGCCAAACACTTTAGGTGTAAGTAGTGAAGCCACA GATGTACTTACACCTAAAGTGTTTGGCTGCCTACTGCCTCGG |
| Mouse sh4-ZMYND8 R | AATTCCGAGGCAGTAGGCAGCCAAACACTTTAGGTGTAAGTACATCTGTGGCTTCACTACTTACAC CTAAAGTGTTTGGCCGCTCACTGTCAACAGCAATATACCTTC |

C

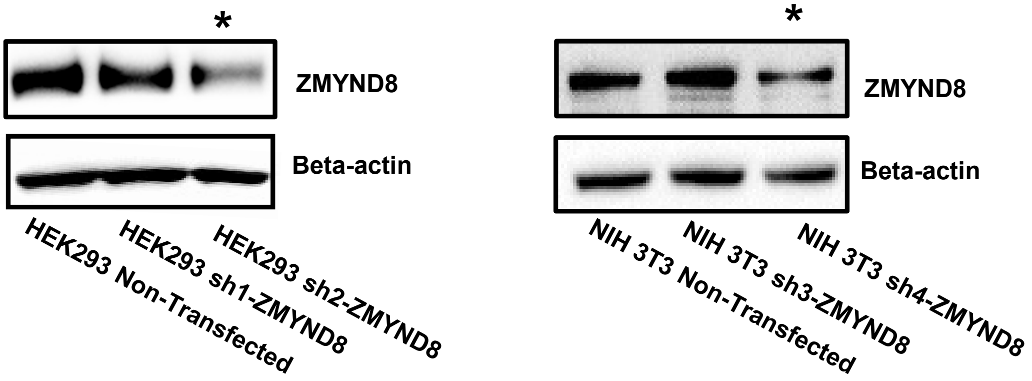

**Figure S4: Generation of short-hair pin (shRNA) knockdown plasmids targeting human and mouse ZMYND8 sequences cloned into the pT2-backbone vector.**
(**A**) Diagram of the plasmids utilized to knockdown expression of ZMYND8 in human (pT2-shZMYND8-hu-GFP) or mouse (pT2-shZMYND8-mo-GFP) cells. (**B**) Corresponding shRNA sequences highlighted in red for sh2-ZMYND8 (human) and sh4-ZMYND8 (mouse). (**C**) Western blot assessing ZMYND8 expression in human HEK293 and mouse NIH3T3 cells 3 days after jet-prime transfection with ZMYND8 knockdown plasmid. Asterisk denotes the shZMYND8 plasmids selected.

**Supplementary Fig S5**

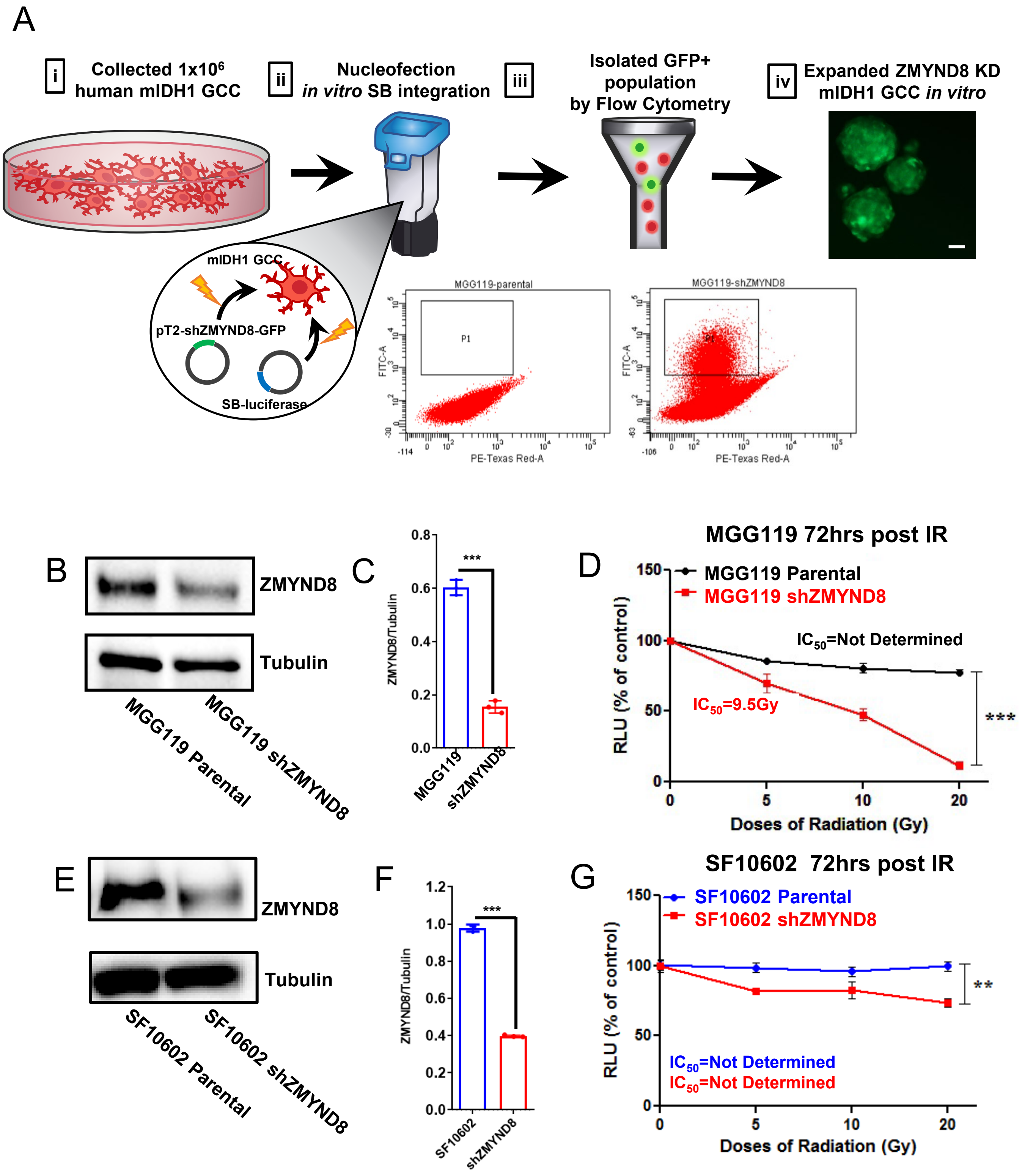

**Figure S5: Generation of shZMYND8 Knockdown mIDH1 GCCs**

**(A)** Illustration of the nucleofection procedure to generate stable shZMYND8 knockdown clones of the human mIDH1 GCCs (SF10602, MGG119). i) $1x10^6$ human mIDH1 GCCs were collected. ii) mIDH1 GCCs were suspended in nucleofection solution containing pT2-shZMYND8-GFP (knockdown plasmid) and sleeping beauty luciferase (SB-luciferase) plasmid to integrate shZMYND8 knockdown plasmid using Lonza nucleofector system. iii) Isolated green fluorescent protein expressing cells (GFP+) by flow cytometry. iv) expanded the stable shZMYND8 cells. (**B**) Western blot for ZMYND8 expression in MGG119 non-transfected (parental) vs. shZMYND8 and (**C**) histogram quantification of ZMYND8 relative to Tubulin. (**D**) CellTiter-Glo assay to assess cellular viability 72hrs post-IR comparing MGG119 parental (black) vs. MGG119 shZMYND8 (red). (**E**) Western blot for ZMYND8 expression in SF10602 non-transfected (parental) vs. shZMYND8 and (**F**) histogram quantification of ZMYND8 relative to Tubulin. (**G**) Cellular viability 72hrs post-IR in SF10602 parental vs. shZMYND8. Errors bars represent standard error of mean (SEM) from independent biological replicates (n=3). ** P<0.01, *** P<0.001; unpaired t test

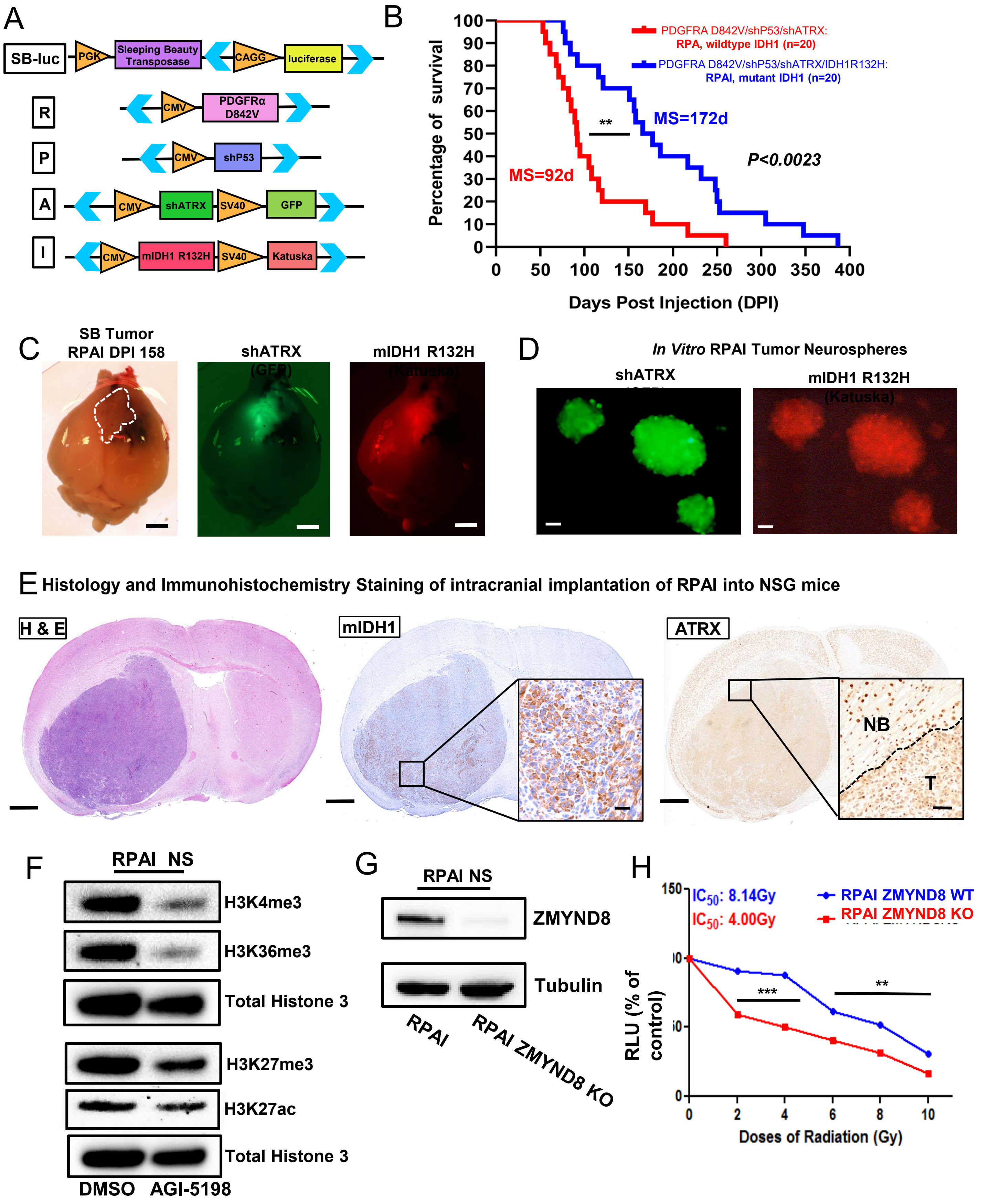
A
SB-luc
PGK
Sleeping Beauty Transposase
CAGG
luciferase
R
CMV
PDGFRα D842V
P
CMV
shP53
A
CMV
shATRX
SV40
GFP
I
CMV
mIDH1 R132H
SV40
Katuska
B
PDGFRA D842V/shP53/shATRX: RPA, wildtype IDH1 (n=20)
PDGFRA D842V/shP53/shATRX/IDH1R132H: RPAI, mutant IDH1 (n=20)
MS=172d
MS=92d
**
P<0.0023
Percentage of survival
Days Post Injection (DPI)
C
SB Tumor RPAI DPI 158
shATRX
mIDH1 R132H
D
In Vitro RPAI Tumor Neurospheres
shATRX
mIDH1 R132H
E Histology and Immunohistochemistry Staining of intracranial implantation of RPAI into NSG mice
H & E
mIDH1
ATRX
NB
T
F
RPAI NS
H3K4me3
H3K36me3
Total Histone 3
H3K27me3
H3K27ac
Total Histone 3
DMSO AGI-5198
G
RPAI NS
ZMYND8
Tubulin
RPAI
RPAI ZMYND8 KO
H
IC50: 8.14Gy
IC50: 4.00Gy
RPAI ZMYND8 WT
RPAI ZMYND8 KO
***
**
RLU (% of control)
Doses of Radiation (Gy)

**Figure S6: Generation of PDGFRαD842V-driven mouse glioma model using SB transposase system.**

(**A**) Schematic representation of sleeping beauty transposase (SB) luciferase (luc) and oncogenic DNA plasmids used to develop gliomas in mice. Blue arrows indicate position of inverted and direct repeat sequences recognized by SB for integration, which flank oncogenic DNA sequences that encode for constitutively active PDGFRA D842V receptor, shP53, shATRX-GFP and mIDH1 R132H-Katuska [RPAI]. (**B**) Kaplan-Meier survival curves for mice bearing wtIDH1 (RPA, n=20) or mIDH1 (RPAI, n=20) gliomas (***P <0.01*, Mantel-Cox test). (**C**) Representative mouse brain of SB RPAI tumor (dotted outline)158 days post injection (DPI) with fluorescence microscopic mages of GFP corresponding to shATRX-GFP plasmid integration and Katuska (red) fluorescence corresponding to mIDH1-R132H. Scale bar: 2mm. (**D**) Fluorescence microscopic images of *in vitro* tumor neurospheres generated from SB-tumor. Scale bar is 500µm. (**E**) Hematoxylin and eosin (H&E) histological stain of NSG mouse brain tumor which was intracranially implanted with RPAI NS. Brain tissue was stained by IHC for mIDH1 and ATRX. Brain section scale bar: 1mm, mIDH1 IHC scale bar: 20µm, ATRX IHC scale bar: 50µm with the tumor boundary defined by dotted line, NB-normal brain, T-tumor tissue. (**F**) Western blot representative of changes in histone mark (H3K4me3, H3K36me3, H3K27me3, H3K27ac) protein expression in RPAI NS treated with either DMSO or AGI-5198 for 1 week. (**G**) Western blot of ZMYND8 expression in RPAI vs. RPAI ZMYND8 KO and (H) Cellular viability 72hrs post-IR in RPAI vs. RPAI ZMYND8 KO. Errors bars represent standard error of mean (SEM) from independent biological replicates (n=3). ** P<0.01, *** P<0.001; unpaired t test.

**Supplementary Fig S7**

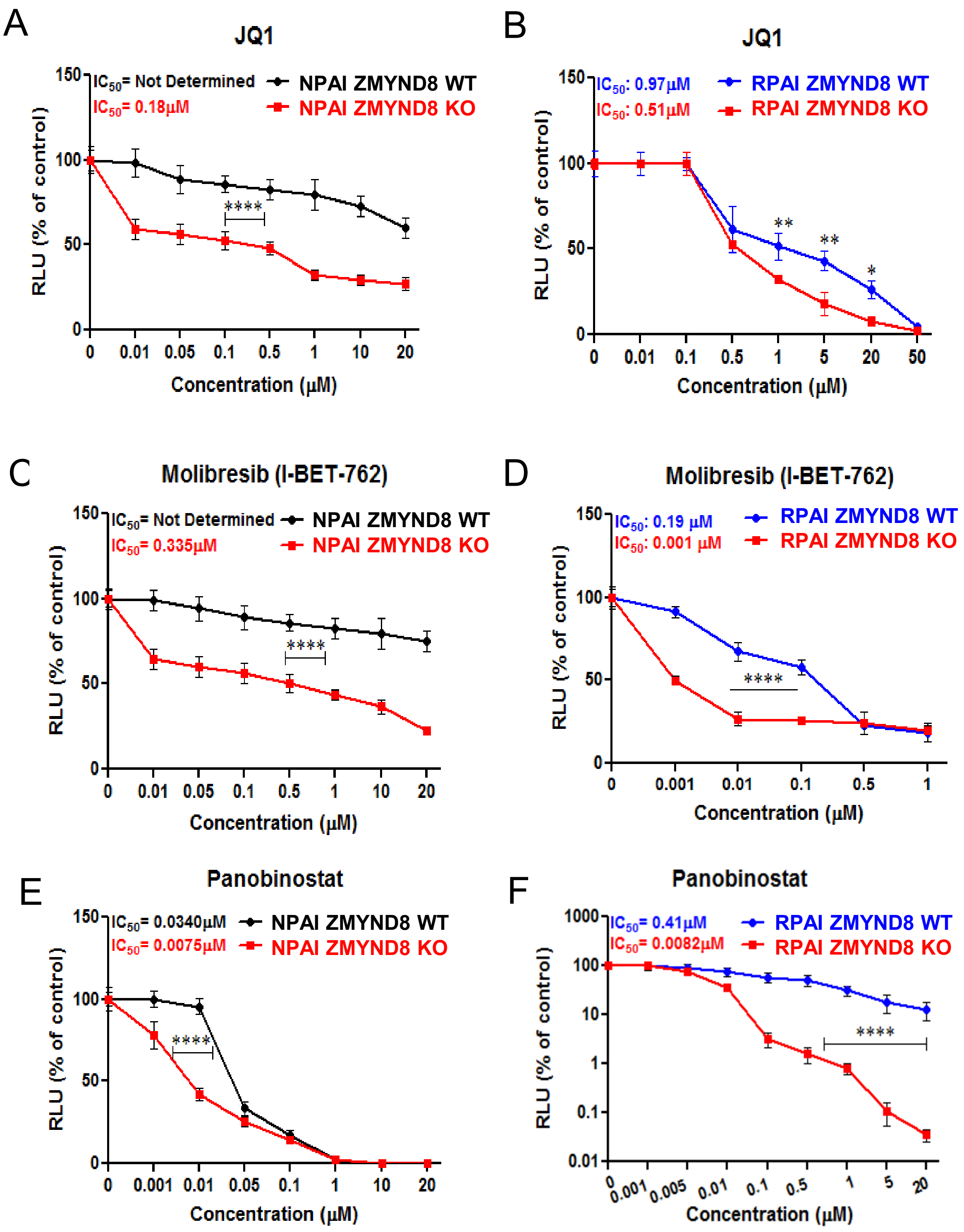

**Figure S7: ZMYND8 KO mouse mIDH1 NS display reduced viability in response to epigenetic inhibitors targeting BRD4 and HDAC compared to ZMYND8 WT NS.**
The impact of epigenetic inhibitors targeting BRD4 (JQ1, I-BET-762) and HDAC (Panobinostat) on *in vitro* cellular viability was assessed in two mIDH1 mouse NS comparing parental (ZMYND8 WT) vs. ZMYND8 KO isogenic clones. Cellular viability was determined based on CellTiter-Glo Assay measured in relative luminescence unit (RLU) compared to non-treated controls. (**A**) Mouse NS harboring NRasG12V, shP53, shATRX, mIDH1R132H mutations [NPAI] ZMYND8 WT (black) vs. ZMYND8 KO (red) treated with JQ1 for 72hrs. (**B**) Mouse NS harboring PDGFRαD842V, shP53, shATRX, mIDH1R132H mutations [RPAI] NS expressing ZMYND8 WT (blue) vs. ZMYND8 KO (red) treated with JQ1 for 72hrs. (**C**) Cellular viability of NPAI ZMYND8 WT vs. ZMYND8 KO treated with I-BET-762 for 72hrs. **(D**) Cellular viability of RPAI ZMYND8 WT vs. ZMYND8 KO treated with I-BET-762 for 72hrs. (**E**) Cellular viability of NPAI ZMYND8 WT vs. ZMYND8 KO treated with Panobinostat for 72hr. (**F**) Cellular viability of RPAI ZMYND8 WT vs. ZMYND8 KO treated with Panobinostat for 72hrs with RLU represented in log scale. The data are representative of three independent biological replicates and error bars denote the SEM of samples performed in triplicate. *P<0.05, ** P<0.01, *** P<0.001, ****P<0.0001; unpaired t test

**Supplementary Fig S8**

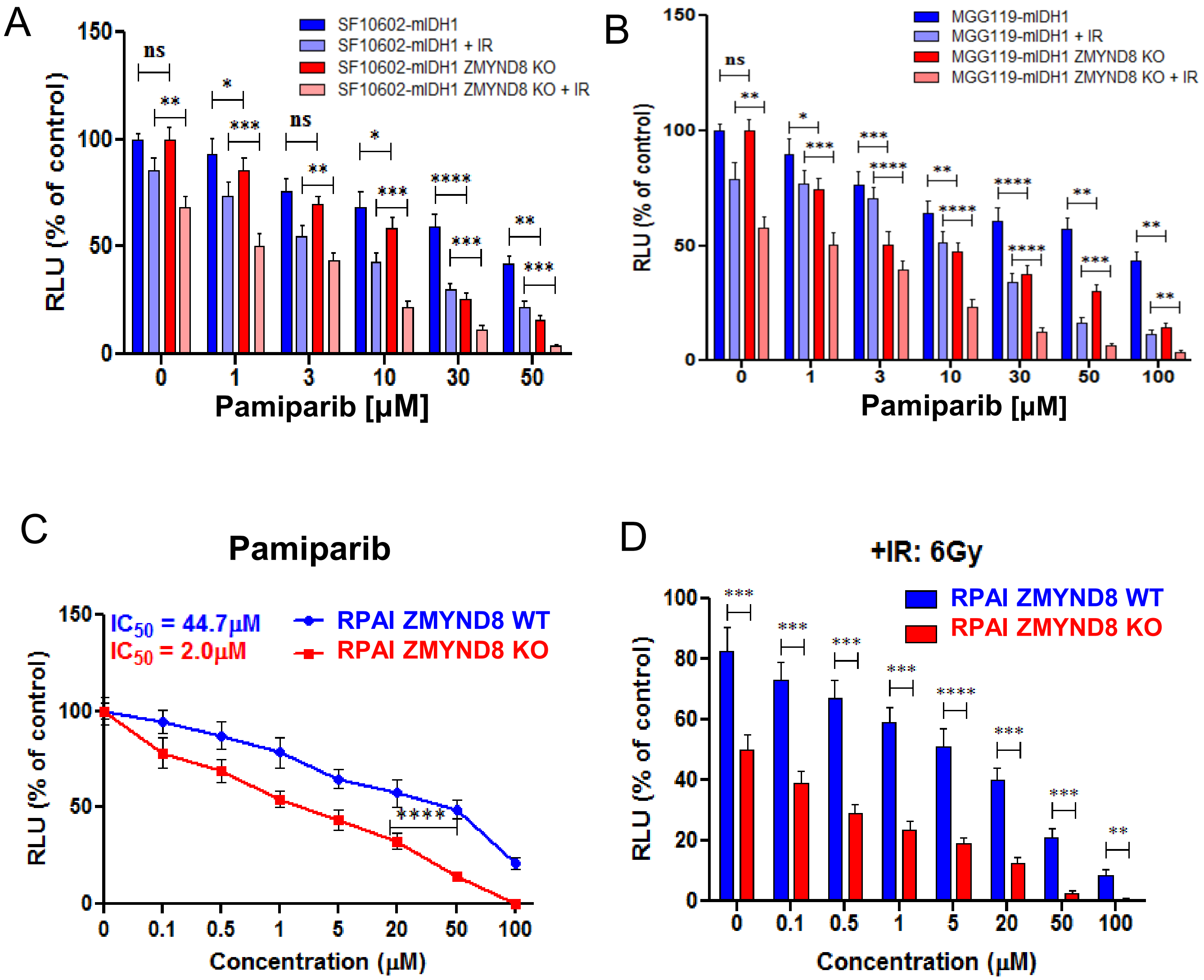

**Figure S8: ZMYND8 KO human mIDH1 GCCs and mouse mIDH1 NS present with reduced cellular viability to IR and in combination with Parp inhibition (Pamiparib).** Representative histogram displays the effect of increased doses of Pamiparib (PARP inhibitor) alone or in combination with IR on cellular viability after 72hrs between (**A**) SF10602 ZMYND8 WT (dark blue) vs. SF10602 ZMYND8 WT + IR (light blue) vs SF10602 ZMYND8 KO (red) vs. SF10602 ZMYND8 KO +IR (pink) and (**B**) MGG119 ZMYND8 WT (dark blue) vs. MGG119 ZMYND8 WT + IR (light blue) vs. MGG119 ZMYND8 KO (dark red) vs. MGG119 ZMYND8 KO +IR (light red). Errors bars represent SEM from independent biological replicates (n=3). Not significant (ns), * $P<0.05$, ** $P<0.01$, *** $P<0.001$, **** $P < 0.0001$; Multiple t test (**C**) Pamiparib dose response curve in RPAI ZMYND8 WT (blue) vs. RPAI ZMYND8 KO (red) to assess cellular viability measured by RLU relative to non-treated control. (**D**) Cell viability assay shows the effect of Pamiparib + IR 6Gy on cell proliferation in RPAI ZMYND8 WT vs. RPAI ZMYND8 KO measured in RLU relative to control non-treated. Errors bars represent SEM from independent biological replicates (n=3) ** $P<0.01$, *** $P<0.001$, **** $P < 0.0001$; unpaired t test

#### Supplementary Fig S9

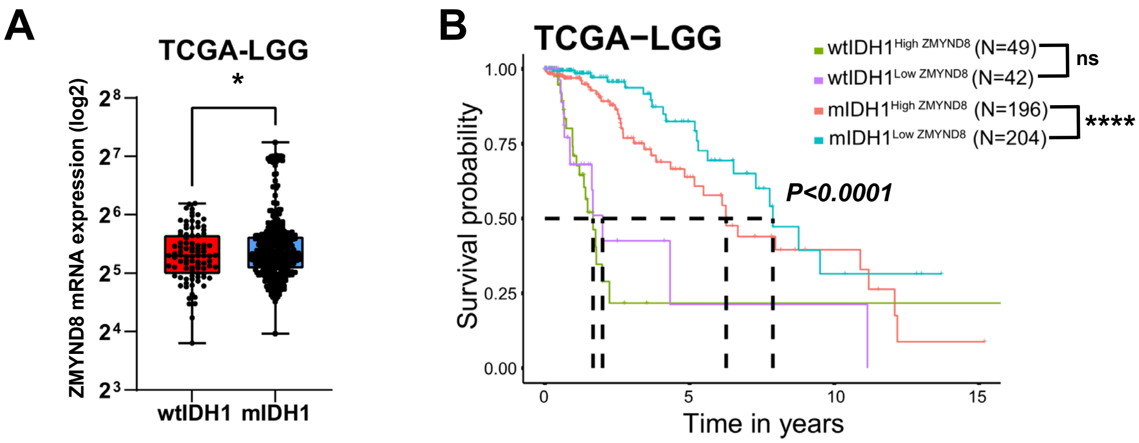

**Figure S9: ZMYND8 is overexpressed in a subset of mIDH1 glioma that is associated with poor clinical outcome in these patients.**
**(A)** Analysis of normalized log2 ZMYND8 mRNA expression levels in human Lower Grade Glioma (LGG) patients from The Cancer Genome Analysis (TCGA) LGG dataset segregated based on IDH1 mutational status as either wildtype IDH1 (wtIDH1) or mutant IDH1 (mIDH1). * $P<0.05$, unpaired t-test. **(B)** Kaplan-Meier survival analysis using the log-rank test for TCGA LGG patients for whom IDH1 mutational status and prognosis data were available. Patients were subdivided by median expression level of ZMYND8 and IDH1 status: wtIDH1 high ZMYND8 (green), wtIDH1 low ZMYND8 (purple), mIDH1 high ZMYND8 (red) and mIDH1 low ZMYND8 (blue). **** $P <0.0001$; log-rank test.

#### Supplementary Table

Table S1. RNA sequencing comparison data

| Sample ID | Description | Number of fragments (2 reads per side) | Number of aligned reads | Percent reads aligned | aligned reads percent duplicates | aligned reads not multi- mapped or discordant |
|---|---|---|---|---|---|---|
| Sample_130939 | Untreated SF10602 rep1 | 54,467,387 | 60,255,501 | 55 | 25.18 | 27,582,011 |
| Sample_130940 | Untreated SF10602 rep2 | 49,159,298 | 48,925,554 | 50 | 23.68 | 22,431,351 |
| Sample_130941 | Untreated SF10602 rep3 | 51,769,590 | 58,220,882 | 56 | 25.39 | 26,813,994 |
| Sample_130942 | Mutant IDH1 inhibitor rep1 | 49,790,214 | 53,857,970 | 54 | 25.23 | 24,706,207 |
| Sample_130943 | Mutant IDH1 inhibitor rep2 | 52,332,347 | 58,386,172 | 56 | 25.92 | 26,662,389 |
| Sample_130944 | Mutant IDH1 inhibitor rep3 | 41,863,074 | 48,099,917 | 57 | 24.67 | 21,962,991 |
